# Accessing the specific capacity of TIL-derived CD8 T-cells to suppress tumor recurrence in resectable HBV-HCC patients

**DOI:** 10.1101/2024.12.05.626571

**Authors:** Janine Kah, Lisa Staffeldt, Gregor Mattert, Tassilo Volz, Kornelius Schulze, Asmus Heumann, Maximillian Voß, Marie-Charlotte Hoell, Meike Goebel, Sven Peine, Maura Dandri, Stefan Lüth

## Abstract

**Background:** Hepatocellular carcinoma represents a significant global health challenge, affecting over a million patients annually, arising mainly from chronic liver diseases, with a majority being related to viral infections. However, despite the groundbreaking clinical results of immune checkpoint blockades and adoptive cell therapies, we still face non-responders accompanied by high rebound rates after resection. Considering that the main concern is to overcome a highly specialized immunosuppressive tumor microenvironment of the individual patient, the characterization of the particular tumor microenvironment and the source of the immune cells used in ACT is of immense importance. Approved ACT therapies mainly use modified peripheral blood cells from individuals. At the same time, tumor-infiltrating lymphocytes are underrepresented even if they have garnered interest due to their potential to target tumor-specific antigens more effectively.

**Methods:** In this study, we employed allogenic and autologous immune cell sources for expansion and stimulation, resulting in adoptive T-cell transfer experiments determining the effector cell differentiation and the related anti-tumor effects by the possible implementation of re-stimulation.

**Results:** We determined a high success rate in expanding and stimulating tumor-infiltrating lymphocytes with consistent CD8 T-cell fractions from HCC patients. To showcase the effectiveness of stimulated T-cells from different sources, we generated cell lines derived from the margin and center of an HBV-induced HCC with a highly immune-suppressive TME. We found effective immune responses supported by cell death induction, ferroptosis, proptosis and apoptosis triggered by all sources of T-cells depending on the area of derived tumor cells.

**Conclusion:** Effector T-cell fractions derived from tumor-infiltrating lymphocytes present a viable source for cell-based immune therapy combined with immune checkpoint inhibitors in HCC patients, especially after resection, to suppress rebound strategies of the parental tumor on an individual level.

## Introduction

Hepatocellular carcinoma (HCC) represents an increasing global health challenge, with more than 1 million patients affected annually [1]. In most cases, HCC develops due to chronic liver diseases arising from viral and non-viral risk factors. Despite constant advances in conventional HCC treatment strategies, the high rates of postsurgical recurrence and resistance to chemotherapy or molecular-targeted drugs necessitate the development of other effective therapeutic approaches [2]. Therefore, immune-based strategies were highly interesting as potentially effective treatment pillars for HCC patients [3]. Currently, the immune-based strategy for HCC patients include oncolytic viruses, adoptive cell transfer (ACT) therapy, immune checkpoint inhibitor therapy, and tumor vaccines [4]. While ACT therapy and OVs target and eliminate tumor cells directly, others, including ICI therapy and tumor vaccines, act as modulators and boost the host’s immune system. Although all these approaches are at the forefront of tumor research and treatment, several pitfalls restrict their comprehensive and effective application for the complete eradication of HCC and related metastasis, due to inherent tolerogenicity of the liver and the immunosuppressive tumor microenvironment (TME) [5].

However, following the clinical success in haematological magnalities, ACT is leading to a high interest in approving individual therapy options for solid tumors, like HCC. The difference between ACT approaches lies in T-cell modification in vitro: CAR-T and TCR genetically modified T-cells are genetically engineered to incorporate modified membrane receptors with a high affinity for selected tumor antigens. Serval promising targets have been identified for genetic modification, like GPC3, AFP, NKG2DL, MUC1, CD147, c-MET and the HBV surface protein [6–11]. In HBV-related HCC, recently conducted and ongoing clinical trials demonstrated acceptable tolerability in patients in advanced stages, unsuitable for liver transplantation when performing ACT with T-cells modified to express HBV-specific TCRs [12]. However, CAR-T and TCR-T-cell therapies harbor inherent limitations for treating patients with HCC, which may explain the significantly worse clinical outcomes than those obtained with CAR-T therapy in hematologic malignancies. Antigens targeted by CAR-T or TCR-T-cells in solid tumors are not specific to tumor cells. They can also be expressed on non-malignant tissues, leading to potential on-target-of-tumor toxicity. These effects are usually dose-limiting and include adverse events like cytokine release syndrome and immune effector cell-associated neurotoxicity syndrome. Significant intra-tumoral heterogeneity in neo-epitope expression and clonal expansion of the adaptive immune system in distant regions was also demonstrated for HCC, including HBV-related HCCs [13]. To overcome the peripheral blood-derived T-cells (PBTs) related limitation, employing tumor-infiltrating lymphocytes-derived T-cells (TITs) as a promising alternative but underrepresented strategy for HCC treatment [3]. However, in 2024, the US Food and Drug Administration approved the first TIL-based ACT for unresectable or metastatic melanoma patients. It reached a milestone in TIL therapy for solid tumors. It involves harvesting TILs from a patient’s tumors, expanding them into bioreactors and reinfusing them [14]. TILs possess natural specificity for tumor antigens due to their direct isolation from the tumor microenvironment, allowing them to recognise a broader range of tumor-specific targets and enhancing their efficacy. At the same time, PBTs typically require extensive activation and expansion to attain similar levels of antitumor activity [15, 16]. Furthermore, TILs have been shown to be better equipped to survive the immunosuppressive tumor microenvironment. They can adapt to the unique demands of fighting tumor. In contrast, PBTs may encounter functional limitations and increased risks of off-target effects, making TILs a more promising alternative for solid tumor immunotherapy [17], especially for patients which underwent resection.

To investigate the ability of TILs, to serve as an alternative strategy to escape reoccurrence caused by resistance to commonly used therapies, we employed generated cell lines derived from the margin and the center region to investigate autologous PBTs and corresponding TITs. As proof of concept, we employed allogenic PBTs derived from a healthy donor for ACT. Since TIL-based ACT has its challenges in isolation and enrichment, we first established a workflow, including collecting PBMCs, tissue and subsequently TILs from HCC patients, with two patients representing HBV DNA in the serum. We used material from one HBV-HCC patient to generate LC11R (margin) and LC11Z (center) cell lines, which were subsequently used in the study after the present immune suppression profile was characterised.

## Methods

### Sample collection, purification and isolation

Human blood samples and HCC-derived cells were collected, isolated, preserved and ethically approved as described previously [18]. The collective of healthy primary hepatocytes were obtained after isolation of 4 donor liver tissues which were not used for transplantation: TM14 (female, 52years), TM47 (female, 11 month), TM46 (male, 2month), TM52 (female, 30years). The study was approved by the Ethical Review Committee of the Ärztekammer Hamburg (WF-021/11). Handling of the human material was performed in accordance with national guidelines and the 1975 Declaration of Helsinki [19].

### Generation of patient-derived Cell lines

Patient-derived primary like cell lines LC11R and LC11Z have been generated and maintained as described previously [18] using orthoptic transplantation using transgenic NOD.Cg.*Prkdcscid Il2rgtm1Wjl/SzJ* background (Strain #:005557) mice.

### Flow cytometry of isolated HCC-cells

Isolated human HCC-cells derived from murine livers after injection of primary tumor cells were characterised as described previously [18], by using the following monoclonal antibodies: anti-CD13, anti-CD133-1, anti-CD24, anti-epithelial cell adhesion molecule (EpCAM), anti-human leukocyte antigen G (HLA-G), anti-CD90, anti-CD73, anti-CD105, anti-CD44 and anti-CD45, conjugated with phycoerythrin (PE), PE-Vio 615, PE-Vio 770, APC and APC-Vio 770. Cells were stained in 1x DPBS containing 0.5% bovine serum albumin fraction (BSA, pH 7, GE Healthcare, Pasching, GER) and 0.05% sodium azide NaN3 (Sigma, Steinheim, GER). The staining was divided into five antibody panels to avoid overlapping emission spectra. Each panel requires 1×104 cells. Positive cells were distinguished from negative cells using unstained samples. Live/dead staining was performed in advance in 1x DPBS (Gibco, Paisley, GB) using Viability Dye 405/520 Fixable Dye according to the manufacturer’s specifications. FACS measurement was carried out on the MACSQuant® Analyzer 10 Flow Cytometer. All antibodies, Viability Dye, and Flow cytometer were received from Miltenyi Biotech (Bergisch Gladbach, GER).

### Enrichment of patient-derived immune cells

Thawed PBMC and TILs were cultivated on a G-Rex24 Well Plate [20] (Wilson Wolf Manufacturing, Minnesota, USA) using 8 ml CTS™ OpTmizer™ (Gibco, Thermo Fisher, Massachusetts, USA) with 10% human AB Serum, 1% Sodium Pyruvate (100 mM) (Gibco, Thermo Fisher, Massachusetts, USA) and 1% L-Glutamin (200 mM) (Gibco, Thermo Fisher, Massachusetts, USA) per Well. For T-cell activation 400 IU/ml rh IL-2 (Sartorius, Göttingen, Germany) and 25 µl CD3/CD28 Dynabeads™ (Gibco, Thermo Fisher, Massachusetts, USA) were added. The PBMC and TILs were cryopreserved 14 days after enrichment.

### HLA Typing

DNA has been extracted from patient blood cell samples for HLA Class I typing using the DNA Isolation Kit (Wizard HMW DNA Extraction Kit from Promega, GER). DNA was used for Luminex-based high-definition LABType rSSO typing (One Lambda, Canoga Park, CA) of the HLA loci HLA-A, HLA-B and HLA-C. LabType SSO is a reverse SSO DNA typing method (rSSO) with sequence-specific Oligonucleotide probes that specifically bind to homologous sequence sections of certain HLA alleles. Genomic DNA (5-60ng/µl) was PCR-amplified using locus-specific biotinylated-primers (Exons 2,4 and 5 for HLA-A and -B; exons 2,4,5,6 and 7 for HLA-C) and Amplitaq Polymerase (One Lambda, Canoga Park, CA), resulting in biotinylated-amplicons. The presence of biotin in each amplicon was detectable using R-Phycoerythrin-conjugated Streptavidin (SAPE). The PCR product was denatured and hybridised to complementary DNA probes bound to fluorescently coded beads. Samples were measured by a flow analyser (FlexMap 3D®, Luminex®, Austin, TX), which identified the fluorescent intensity of phycoerythrin (PE) on each bead. HLA Fusion^TM^ program Version 4.6.1 (One Lambda, Canoga Park, CA) was used to analyse the data.

### Flow cytometry of Immune cells

Flow cytometry analyses of the therapeutic compartment were performed at D1 and D4 of co-culture with LC11R and LC11Z. Analysis was performed on the SymphonyA3™ system (Becton Dickinson, New Jersey, USA) using the antibody panel S10 and S7 (all from Becton Dickinson, New Jersey, USA), described in **Supplementary Table 1**. The antibodies were incubated for 30 minutes at room temperature, and protected from light. Before the staining, the cells were incubated with Human BD Fc Block^TM^ (Becton Dickinson, New Jersey, USA). The gates were defined using fluorescence minus one (FMO) control.

### Cell culture and spheroid formation and treatment

Immortalized HepG2H1.3 and patient-derived cell lines were maintained as described previously [18]. For 2D culture conditions, cells were seeded into xCELLigence e-plates (Agilent, Santa Clara, United States) for cell index measurement using Agilent’s Real-Time Cell Analysis (RTCA) or into conventional 24-well plates for RNA Isolation and microscopic captures of unstained or stained LC11 cells, aT-cell numbers ranking from 10 - 150000 cells per well. LC11 cells were allowed to growth for 24h before treatment was performed. For 3D conditions, different amounts (5000 – 30000 cell/ well) of LC11R and LC11Z cells were seeded on BIOFLOAT™ cell culture plates in DMEM containing L-glutamine and glucose, supplemented with 1% P/S, 10% Gibco fetal bovine serum (FBS; all from Thermo Fisher Scientific, Waltham, USA) and were obtained for 3 weeks to follow the spheroid formation. Spheroids were used for co-culture experiments after 3 weeks of formation. For visualisation, LC11R cells stably transduced with the vector LeGO-iG2-Puro+-Luc2 (3rd generation HIV1-derived self-inactivating vector) were used. Stable transduction was performed as described previously [18]. For treatment under 2D cell conditions, the supernatant was collected before ACT. For ACT, T-cells derived from HD24:02, HD02:01 or the HBV-HCC patient were thawed and cultivated for 3 days in 6-well plates with recombinant hIL-2 (180IU/ml) and CD3/CD28 beads. After 3 days, stimulated T-cells were washed and counted, and ACT was performed on the indicated target-to-effector cell ratios (T:E). To visualise the transferred T-cells, a cell tracker solution purchased from ThermoFischer Scientific (Waltham, USA) was used in 5µm dissolved in PBS, following the manufacturer’s instructions. During treatment, supernatant and / or T-cells were collected for analysis, and captures were obtained manually or automatically using the BX-780 Microscope (Keyence, Osaka, Japan). For re-stimulation treatment, rhIL-2 was applied to the indicated specimens 24h after ACT start.

### Virological load measurement

Extraction of HBV DNA from serum samples and supernatant was conducted with the QiAmp MinElute™ Virus Spin kit (Qiagen, Hilden, Germany). For quantification, TaqMan PCR was performed using an HBV-specific probe, as listed in **Supplementary Table 2**, and cloned HBV-DNA references were amplified in parallel to establish a standard curve for quantification.

### Isolation of cell-free DNA

Cell-free DNA (cf-DNA) obtained from human plasma or serum was collected, preserved and isolated using the MagMAX™ Cell-Free DNA Isolation Kit (ThermoFisher, Massachusetts, USA). Before isolation, samples were denatured with protein K and SDS (sodium dodecyl sulfate) an incubated for 20 minutes. For the isolation, the samples were mixed with MagMAX™ Cell Free DNA Lysis/Binding Solution and MagMAX™ Cell Free DNA Magnetic Beads, incubated for 20 minutes, then cooled for 5 minutes. After the beads bound to the cf-DNA, the solution was washed successively with MagMAX™ Cell Free DNA Wash Solution and 80% ethanol, centrifuged and the supernatants were removed. The obtained cf-DNA was eluted in the final step with MagMAX™ Cell Free DNA Elution Solution.

### Real-Time PCR measurement

All qPCR reactions were performed using 1 µl of eluted cf-DNA, 7,5 µl PowerUp™ SYBR™ Green Master Mix (ThermoFisher, Massachusetts, USA), 4,5 µl water and 1µl of paired primer, as listed in **Supplementary table 3**. Cf-DNA was quantified by amplification of short and long fragments of the LUC insert, shown in **Supplementary Figure 2A** (LUC primer 1, 2, 4 and 10), Alu 1 and MTCO (1 and 2) markers were targeted with qPCR (QuantStudio™ 7 Pro Real-Time PCR System, Applied Biosystems, Massachusetts, United States). Cycling conditions consisted of initial denaturation at 95°C for 2 min, and 40 cycles of 95°C for 15 s and 60 °C for 1min and cooling at 20°C for 2 min. Analysis was performed using the Design and Analysis Software from Applied Biosystems,

### Measurement of spheroid area

Spheroids were captured at indicated timepoints automatically using the BX-780 Microscope and were analysed by calculating the area using the Marco Cell Count™ module for automatic analysis (Keyence, Osaka, Japan).

### Isolation of Oligonucleotides

RNA was extracted from liver specimens cell lines in 2D and 3D conditions and from co-cultured cells using the RNeasy Mini™ and Micro™ RNA purification kit (Qiagen, Hilden, Germany) [21].

### Measurement of gene expression level using TaqMan-based PCR

For measurement of gene-expression, 2-step PCR was performed. Therefore, complementary DNA (cDNA) synthesis was conducted by using MMLV Reverse Transcriptase™ 1st-Strand cDNA Synthesis Kit (Lucigen, Middleton, Wisconsin, USA) to synthesise RNA complementary DNA, according to the manufacturer’s instructions. Human-specific primers from the TaqMan Gene Expression Assay System, listed in **Supplementary Table 2**, were used to determine gene expression levels (Life Technologies, Carlsbad, California, USA). Samples were analysed using the Quant Studio 7™ Real-Time PCR System (Applied Biosystems, Massachusetts, United States). The human housekeeping gene ribosomal protein L0 (RPL0) was used to normalise human gene expression levels.

### RNA Isolation and Sequencing

Total RNA was extracted from primary hepatocyte samples, patient-derived HCC specimens and primary HCC cell line LC11R using the RNeasy Mini Kit (Qiagen, Hilden, Germany), following the manufacturer’s protocol. The quality and quantity of RNA were assessed using a NanoDrop spectrophotometer (ThermoFisher, Massachusetts, USA) and Agilent Bioanalyzer 2100 (Agilent, Santa Clara, United States). High-quality RNA samples with an RNA Integrity Number (RIN) greater than 7.0 were selected for sequencing.

### Library Preparation and RNA Sequencing

RNA sequencing libraries were prepared using the TruSeq Stranded mRNA Library Prep Kit (Illumina, Californien, USA) according to the manufacturer’s instructions. Briefly, mRNA was purified from total RNA using poly-T oligo-attached magnetic beads and fragmented into small pieces. First-strand cDNA was synthesised using random hexamer primers and reverse transcriptase. This was followed by second-strand cDNA synthesis, end repair, A-tailing, adapter ligation, and PCR amplification to enrich the cDNA fragments. The libraries were quantified using a Qubit fluorometer (Thermo Fisher, Massachusetts, USA) and assessed for size distribution using the Agilent Bioanalyzer 2100. The libraries were then sequenced on the Illumina NextSeq 1000 platform, generating 150 bp paired-end reads. Technical replicates were used for RNAseq for each sample (n=2).

### Data Processing and Differential Expression Analysis

Raw sequencing reads (n = 2 per sample) were prepared as R input data sets and further processed with R [22]. Input was trimmed using Trimmomatic [23]. Clean reads were aligned to the human reference genome (GRCh38) using STAR aligner [24]. The resulting BAM files were processed with featureCounts to generate read count matrices [25]. Differential expression analysis was performed using the DESeq2 package in R [26]. Genes with an adjusted p-value (Benjamini-Hochberg correction) of less than 0.05 and a log2 fold change (log2FC) greater than 2 or less than −2 were considered significantly differentially expressed.

### Data Processing and Visualisation

DEGs were used for volcano plot generation using the DESeq2 package in R [26]. DEGs were implemented for pathway comparison analysis and analysis match with global datasets using the Ingenuity Pathway Analysis™ Software from Qiagen (Hilden, Germany) [27]. Differential regulation of canonical pathways has been extracted by setting p-value > 0.05 log10 and z-scores > 3,5. Genes, which were up- or downregulated in the indicated pathways are colored red or green. The genes overlapping between the comparison analysis and the indicated pathways were numbered at the top of the columns.

### Protein analysis by immunofluorescence

For histological characterisation cultured cells were processed as described previously [18, 28]. Seeded and treated cells were washed with 1x PBS and fixed with 4% PFA for 10 min. After fixation, generated LC11 cells and immortalised cell line HepG2.1.3 were used for immunofluorescence staining against indicated proteins using primary antibodies, rabbit anti-caspase 3 (710431, Invitrogen, 1:100), rabbit anti-PDL1 (GTx57193, GeneTex, 1:100) and rabbit anti-Calnexin (MA5-32332, ThermoFischer; 1:200). Specific signals were visualised with Alexa 555 labelled secondary antibodies (Invitrogen, Darmstadt, GER). Nuclear staining was achieved by Hoechst 33258 (Invitrogen, Eugene, USA). Stained cells were analysed by fluorescence microscopy (BZ-9000 and BX-780, Keyence, Osaka, Japan). Captures of tissue slides were generated manually with consistent exposure times for equal material and antibodies. For LC11, spheroids captures were generated automatically with the same magnifications and exposure time for the same antibodies and fluorescence.

### GZMB quantification

ELISA targeting Granzyme B were conducted using the Human Granzyme B DuoSet ELISA (R&D Systems, Minnesota, USA). The procedure was done according to the manufacturer’s protocol with slight alterations for the Human Granzyme B DuoSet ELISA ELISA Assay Diluent (5X) (Biolegend, California, USA), which was used to block the plates and dilute the reagents. Streptavidin-PolyHRP20 (SDT GmbH, Baesweiler, Germany) was used instead of the peroxidase contained in the Kits. The Streptavidin-PolyHRP20 was incubated for 30 minutes at room temperature on a shaker. Incubation was stopped using a stopping solution, and measurement was conducted using the Tecan Plate Reader Spark® and the corresponding Magellan Software™.

### Cytokine and Chemokine Quantification

The LEGENDplex™ Human Inflammation Panel 1 (13-plex) (BioLegend, San Diego, CA) was used to quantify 13 pro-inflammatory cytokines and chemokines (IL-1β, IFN-α2, IFN-γ, TNF-α, MCP-1, IL-6, IL-8, IL-10, IL-12p70, IL-17A, IL-18, IL-23, IL-33) in cell culture supernatants. Samples were incubated with antibody-conjugated beads, followed by detection with biotinylated antibodies and a Streptavidin-PE label. Data acquisition was performed on the SymphonyA3™ system (Becton Dickinson, New Jersey, USA), and concentrations were calculated using standard dilutions in the LEGENDplex™ Data Analysis Software. For relative quantification, supernatants from untreated controls or cell culture mediums were used.

### Statistics and Sample Sizes

Graph design and statistical analysis software GraphPad Prism Version 9 (GraphPad Software, Inc., La Jolla, CA, USA) was used. Cell index differences were analysed using a two-tailed unpaired t-test of 5 biological replicates for each group within the indicated timepoints (**Figures 2C**, **Figure 3F**, **Figure 5A, B and Figure 6B**). For 2D RNA isolation, n = 3 wells were used and pooled for the lysis. For 3D RNA isolation, n = 3 spheroids from each group were used und pooled for the lysis. For all cDNA analysis using TaqMan assays, n = 3 technical replicates were used. Expression comparison was analysed using a two-tailed unpaired t-test of the 3 technical replicates for re-stimulated against non-re-stimulated groups for 3D cultures (**Figure 3A**, **Figure 6A**, **Figures 7K, N and O – Q**). Area size comparison was performed using a two-tailed unpaired t-test of 3 biological replicates for each group (**Figure 7B**). For Cytometric analysis, samples from biological replicates were pooled and divided into n = 2 replicates for each staining panel (**S10, S7 and Legendplex multiplex analysis**). Panels were measured and analysed separately. Statistical outputs are indicated in the figure legends; p-values were plotted in the graph as follows: * p < 0.05; ** p ≤ 0.01 and *** p ≤ 0.001, **** p ≤ 0.0001.

## Results

### Isolation and expansion of Tumor-infiltrating lymphocytes result in consistent CD8 T-cell enrichment

Immune cells obtained from the center of the parental tumor and the blood were utilized for expansion, as shown in **Figure 1A**. Immune cells from six patients with advanced HCCs were employed for the preliminary isolation and expansion. Immune fractions were examined for the lineage marker expression during the scaling process, as illustrated in **Figure 1B**. Two patients were clinically identified as HBV DNA serum positive on the day of resection. These patients are indicated with red and purple dots in **Figure 1B**.

**Figure 1:**
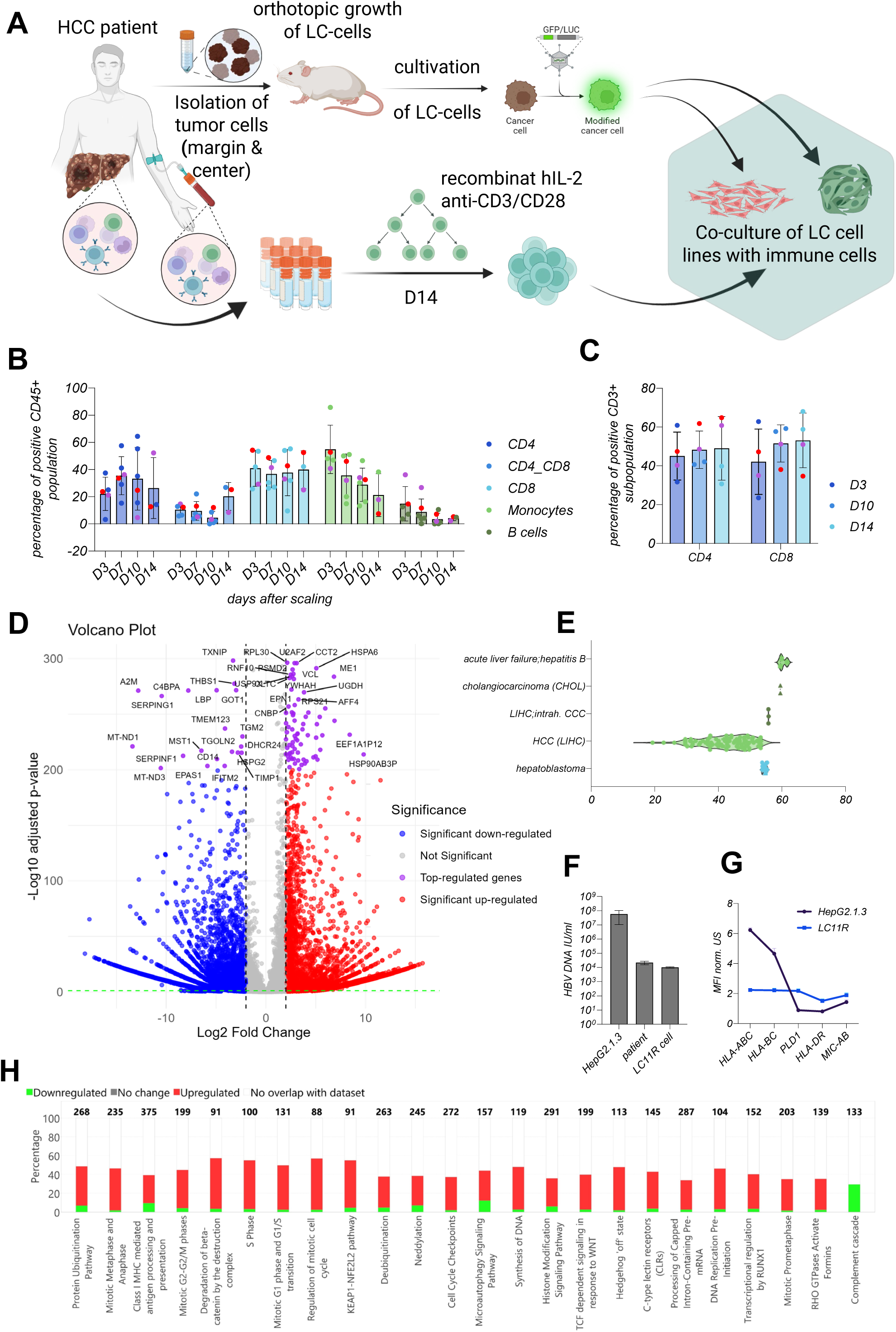
Generation of patient-derived cell lines and expansion of corresponding immune cells. In **Figure 1A**, the experimental setting is illustrated. Material from HCC patients undergoing resection were isolated and used to generate cell lines derived from the margin and the center region. From the same patients, blood and tissue-derived immune cells were isolated. All isolated cell types were cyro-conserved. For the expansion of immune cells from blood and tissue, 6 patients were employed and observed over time, as depicted, regarding the percental distribution of CD45+ (**Figure 1B**) and CD3+ (**Figure 1C**) subpopulations. Cell types were plotted as indicated, and HBV-related HCCs were plotted in purple and red for individual tracking. The generated margin-derived LC11R cell line was characterized and classified in D-H using DEG and DEG-based pathway analysis. The volcano plot illustrates the differential expressed genes relative to healthy hepatocytes (PHH), with indicated significances. Top-regulates genes are plotted in purple and marked with ensemble names. In **Figure 1E**, canonical and disease-related pathways were matched to the global analysis. High z-scores indicate high similarity in pathway prediction scores. Here, all positive scores were sub-grouped and plotted using a violin plot. In **Figure 1F**, the HBV DNA titers in IU/ml were measured using TaqMan PCR. They quantified using a high HBV control diluted standard in the serum of the individual patient used in the study, the HBV-producing cell line HepG2.1.3 and the supernatant of the LC11R cells after generation. In **G**, the presence of depicted molecules analyzed by flow cytometry are plotted, using the HepG2.1.3 cell line as a control line to classify the margin LC11R cells regarding antigen presentation. In **Figure 1H**, canonical pathways with z-scores > 3,5 and p-values > 0,05 were extracted. Gene numbers relevant to the individual pathway and measured in DEG analysis of LC11R against PHHs, were mentioned on the top of the columns. Downregulated genes were marked in green, and upregulated genes were marked in red.

In general, monocytes and B cells exhibited a persistent decline during expansion. The distribution was very heterogeneous for CD4- and CD4/CD8-positive T-cells, whereas CD8 T-cells constituted a stable fraction during expansion. The patient depicted with red dots showed reduced CD4 and elevated CD4/CD8 titers, accompanied by a stable CD8 T-cell titer following expansion compared to the baseline sample. Conversely, elevated CD4 and diminished CD8 titers for the same patient were observed upon expansion for the blood-derived immune cells, as illustrated in **Figure 1C**. However, both fractions were cryopreserved after 14 days of expansion. Thawed tissue-derived and blood-derived immune cell compartments from day 14 were utilized for the adoptive T-cell transfer to examine the influence of the immunological signature and the origin of the immune cells.

### Generated patient-derived margin cell line LC11R mirroring immune evasion characteristics of the parental malignancy

To investigate the therapeutic effectivity of re-stimulated immune cells obtained from blood and tissue of the HBV-infected patient, within their autologous tumor microenvironment, patient-derived tumor cell lines were generated from the margin and center materials, as described previously [18] and shown in **Figure 1A**. Differential gene expression analysis compared to healthy liver cells was conducted to characterize LC11R cells, as illustrated in **Figure 1D**. Here, biochemical alterations delineate the transition from healthy tissue to a malignant phenotype. Genes linked to cellular stress response and proliferation in hepatic tumor cells were activated significantly. Heat shock proteins (e.g., HSPA6, HSPA4) significantly increase. The increased concentrations of these proteins indicate an enhanced cellular defense response to stress, potentially allowing tumor cells to endure and adjust to the adverse microenvironment typically found in tumors. Alongside stress response genes, other markers associated with cell cycle control and proliferation were increased, including ribosomal proteins (e.g., RPL genes) and JAK2. This rise indicates a transition towards augmented cell division and growth signaling pathways often triggered in malignant cells. Genes associated with protein translation and ubiquitin-mediated degradation (e.g., UBE2D3, TRIP12) were also elevated. This pattern indicates a compensatory mechanism in tumor cells that increases protein production and modifies breakdown pathways, promoting tumor cell survival and proliferation.

In contrast, numerous essential genes exhibited marked downregulation in hepatic tumor cells, especially those associated with inflammatory responses and death. Prominent among these were genes like CD14 and THBS1, which mediate the immune response. The diminished expression may indicate a strategic adaptation by tumor cells to avoid immune identification, improving their survival chances within the host. Metabolic genes, such as TGM2 and LBP, were downregulated, indicating modifications in metabolic pathways typically linked to tumor. This transition enables a metabolic reconfiguration towards the Warburg effect, in which tumor cells predominantly rely on glycolysis for energy generation, despite oxygen availability. This metabolic adaption facilitates fast growth and modifies the tumor microenvironment.

Moreover, apoptosis and neuroprotection-related factors were downregulated, including SERPINF1 and IFITM2. This reduction indicates resistance to apoptosis, facilitating the increased survival of tumor cells despite genetic instability. The findings highlight the pathway modifications, illustrated in **Figure 1H**, that characterize the carcinogenesis of the examined LC11R cells. The match analysis illustrated in **Figure 1E** depicts the classification of the generated LC11R cells by a comprehensive global comparison. The match analysis to global data sets, shown by elevated z-scores concerning canonical and disease pathways regulation, was attained with HCC-related datasets, as illustrated by violin plots.

Furthermore, the highest accordance was observed in HBV-related HCCs and acute liver failure, highlighting the representative characteristics of the produced LC11R cell line. **Supplementary Figure 1** illustrates the surface proteins on LC11R cells. The expression of CD133, CD13, CD90, and CD326 signifies a tumor stem cell-like population that enhances tumor heterogeneity, therapeutic resistance, and the aggressiveness of liver tumor. The lack of CD34, CD24, CD44, and CD105 indicates a differentiation status of the LC11R cells that will affect their behavior and treatment responses. Compared to immortalized HBV-producing liver tumor cells (HepG2.1.3), MIC-AB, PDL-1, and HLA-DR expression were elevated. At the same time, MHC class I proteins were expressed at reduced levels in LC11R cells, as illustrated in **Figure 1F**. The results demonstrate the retention of the immune evasive characteristics from the original HCC in the developed cell line. **Figure 1G** reflects the detection of HBV DNA in the patient’s serum and the supernatant of cultivated cells, compared to genetically modified HepG2.1.3 cells. LC11R cells produced HBV DNA at low titers comparable with the parental production.

### Allogenic PBMC-derived CD8 T-cells exhibit sustained cell death induction in parental margin-derived cells

Initially, CD3/CD28-enriched PBTs from healthy donors (HD24:02 and HD02:01) with an anti-HBs titer > 100 IU/L were used for the adoptive T-cell transfer. HD24:02 provided the matched PBTs for LC11 cells, while HD02:01 provided matched PBTs for the immortalized cell line HepG2.1.3 (see Suppl. Fig. 3). The HD-PBT-ACT was additionally conducted for GFP/LUC-producing LC11R_LUC cells, as shown in **Figure 2A** and described in **Supplementary Figure 2**. In **Figure 2A**, the experimental settings are illustrated. **Figure 2B** demonstrates that the induction of apoptosis following ACT of HD24:02 was assessed post-treatment. A pronounced cytolytic effect, illustrated in **Figure 2C**, through the assessment of cell proliferation, was determined over time. In contrast, HD02:01 PBTs exhibited ineffective lysis of the LC11R cells, but produced GZMB in high titers, as shown in the **right panel of Figure 2C**. HD02:01-PBTs were evaluated under identical conditions, utilizing HepG2.13 cells to assess the effectivity, as shown in **Supplementary Figure 3**. Here, HD02:01 PBTs successfully lysed HepG2.1.3 cells, as shown in the cell index measurement (**Suppl. Fig. 3B**) and the live imaging over time, employing low (**Suppl. Fig. 3C**) and high (**Suppl. Fig. 3D**) initial cell numbers for ACT.

**Figure 2:**
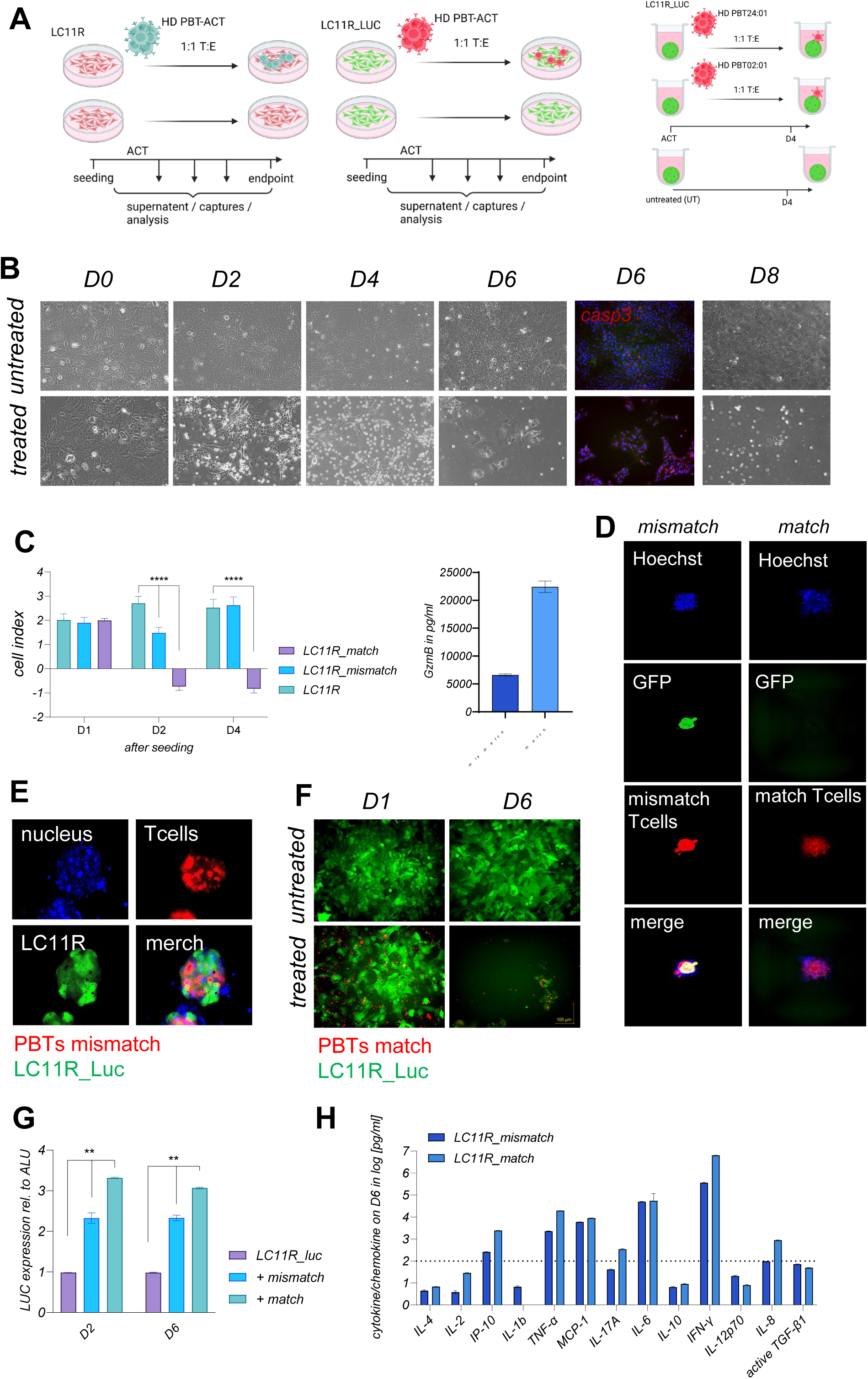
Adoptive T cell transfer with expanded and stimulated healthy donor PBT in immune suppressive and evasive LC11R cells. In **Figure 2A** the experimental setting for the proof-of-concept ACT is shown, using expanded and stimulated PBT from healthy donors, presenting HLA-A 24:02 or HLA-A 02:01, following LC11 or HepG2.1.3 HLA-types. ACT was performed one time either in the use of conventional 2-dimensional (2D) culture (B, C and F-H) or by employing 3-dimensional (3D) culture (**Figures 2D and E**). Following the HLA match of the cell line of interest, HLA-A 24:02 is marked as matched and HLA-A 02:01 as mismatched PBTs. In **Figure 2B**, ACT using matched conditions was followed by automatic microscopy for the same region of interest for 8 days, whereby brightfield captures are shown on the indicated time points and conditions. On day 6 after ACT, the expression of caspase 3 was evaluated by immunofluorescence staining. **Figure 2C** shows the ACT in match and mismatch conditions for cell viability measurement using the xCELLigence system (left) and the GZMB production (right) after 4 days. In **Figure 2D**, LC11R luciferase and green fluorescence protein-producing spheroids are presented after 4 days of ACT using life cell imaging and the conditions shown in **Figure 2A**. Here, PBTs were presented in red, LC11R_Luc in green, and the nucleoli were stained with Hoechst. Single and merged captures were generated automatically. On day 4, spheroids under mismatched conditions were transferred to glass chamber slides, captured in 60xmagification and presented in **E**, using single and merged captures. In F, initial and final captures from life cell imaging of the same regions of interest are shown under 2D conditions for matched PBTs. Co-culture was observed for 6 days. PBTs were marked with a suitable life cell tracker, as described in the methods. In **Figure 2G**, the quantification of cf-DNA by amplification of LUC-specific transcripts, as described in the methods, is shown for the ACT performed for 6 days, whereby supernatant was collected on day 2 and day 6, as indicated. In **Figure 2H**, cytokine and chemokine profile is presented 6 days after transfer, using the same supernatant analyzed in **Figure 2G**. Statistical outputs are p-values were plotted in the graph as follows: * p < 0.05; ** p ≤ 0.01 and *** p ≤ 0.001, **** p ≤ 0.0001.

Nonetheless, HD02:01 could elicit cytotoxicity within the initial hours of co-cultivation, likely attributable to the elevated cytokine generation during the stimulation process. Remarkably, LC11R cells rapidly adapted to this cytokine environment. Until day 3, the cytokine generation by HD02:01 PBTs did not affect proliferation or further lyse the liver tumor cells. Conversely, HD24:02 PBTs inhibited the growth, and additional lyse LC11R cells. This was similarly observed for 3D-based studies conducted under identical conditions for 4 days, as illustrated in **Figure 2D**. **Figure 2E** demonstrates that the mismatched T-cells (depicted in red, labelled with Alexa-555 cell tracker) successfully migrated within the LC11R_LUC spheroids (shown in green) but did not lyse nor disrupt the dense structure.

To quantify the cytolytic effect, LC11R_LUC cells were employed for 2D culture and subsequently HD24:02 PBT-ACT, as preventatively shown in **Figure 2F**. Quantification was done by amplifying released LUC-encoding plasmid DNA (p-DNA) in the cell-free DNA fraction using specific primers (refer to **Supplementary Figure 2A and F**). **Figure 2G** displays the concentration of cell-free LUC p-DNA in the supernatant, which was increased 24 hours after the ACT of PBTs in both conditions.

Nonetheless, the amplified free LUC p-DNA was elevated, corresponding to the findings obtained from cell index measurement. **Figure 2G** and **Figure 2C** illustrate that the lysis peak occurred within 24 hours following ACT. The initial cytokine-induced cell lysis facilitated by HD02:01 PBTs resulted in an increase of LUC p-DNA in the nonspecific co-culture as well. Cytokine and chemokine levels were assessed on day 6 for both conditions, as illustrated in **Figure 2H**. As anticipated, cytokines resulting from specific and nonspecific T-cell responses were elevated. IL-2, IP-10, IFN-γ, TNF-alpha, and IL-8 were produced in more significant quantities in particular samples, but IL-1beta and IL-12p70 were elevated in unspecific samples. The regulatory cytokines IL-17A and IL-10 were present in both conditions, albeit at reduced levels relative to the pro-inflammatory cytokines IP-10, TNF-alpha, IL-6, IL-8, and IFN-γ. Notably, IL-1β, known for its role in the differentiation and proliferation of regulatory T-cells, was only detectable under nonspecific conditions.

Next, both conditions were employed to conduct RNASeq analysis, 48h post ACT. The expressing analysis, shown in **Figure 3A**, displays the significant elevation of HLA-A, B, C and G expression for HD24:02 conditions. A negligible increase was observed in the HD02:01 treated LC11R samples. These findings reinforce the prior conclusion that, under HLA-A matching conditions, T-cells could elicit a specific immunological response, whereas unmatched T-cells did not. After ACT with HD24:02, genes related to immunological functions, apoptosis, DNA repair, and cellular integrity were downregulated, as shown in **Figure 3B**. Conversely, genes associated with tumor development, metastasis, and immune evasion were elevated significantly. The DEG-based pathway analysis, shown in **Figure 3C**, elucidates the induced signaling shift towards immune evasion and diminished anti-tumor immunity, indicating that LC11R cells facilitate immune evasion and increased proliferation in direct response to the ACT.

**Figure 3:**
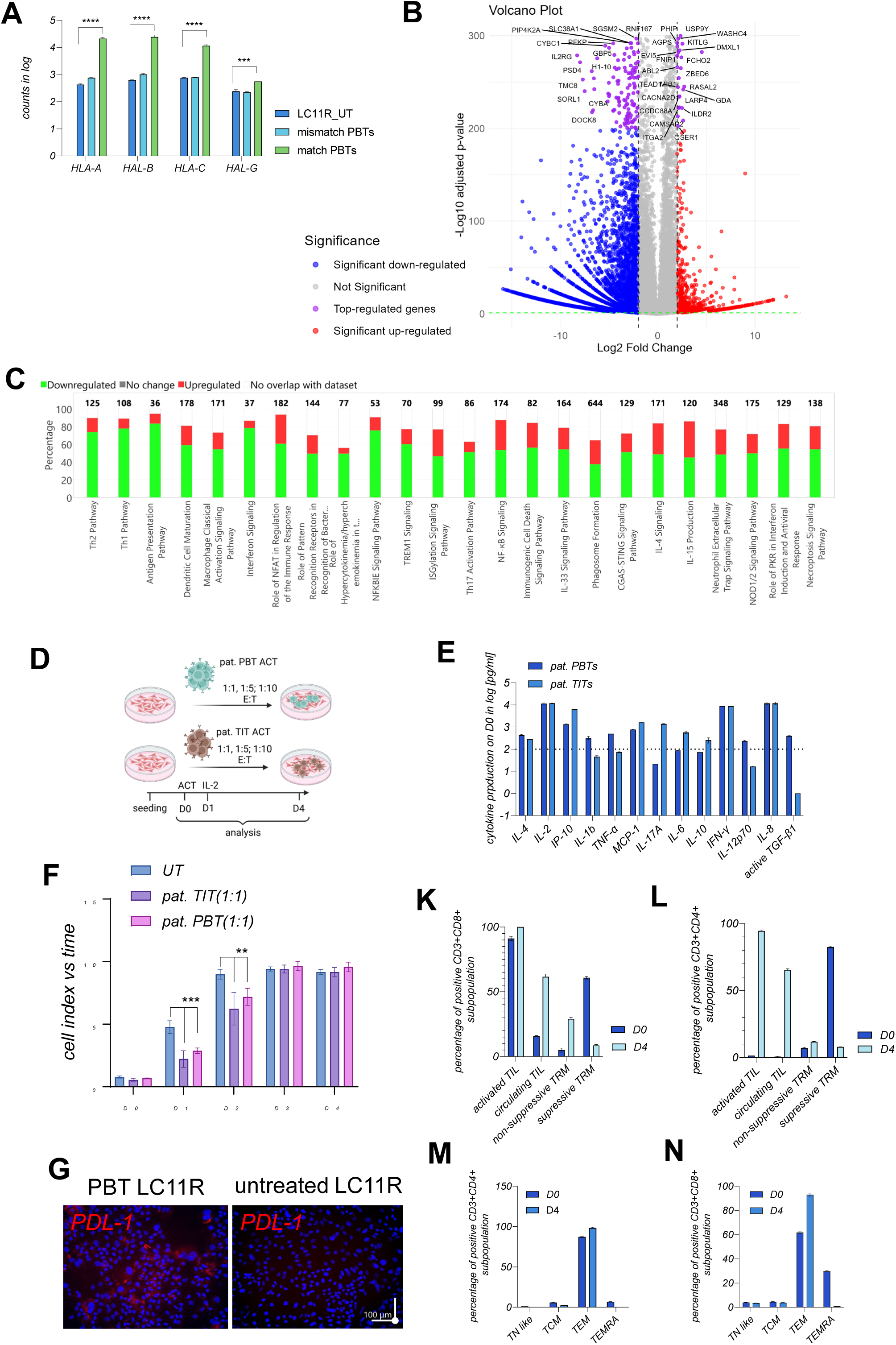
Early timepoint DEG analysis of healthy donor PBTs transferred into LC11R cell culture and the POC-ACT of autologous PBTs and TITs using LC11R cells. The early and prominent cytotoxic effect shown in **Figure 2** was reasonable to extract RNA for DEG at day 2 after ACT. Therefore, in **Figures 3A and B**, the early time point is presented in DEG and pathway analysis (**Fig. 2C**). In **Figure 3A**, the expression of MHC molecules is plotted under the mentioned conditions. In **Figure 3B**, the volcano plot illustrates the differential expressed genes relative to the untreated LC11R cells, with indicated significances. The top-regulator genes are plotted in purple and marked with ensemble names. In **Figure 3C**, canonical pathways with z-scores > 3,5 and p-values > 0,05 were extracted. Gene numbers relevant to the individual pathway and measured in DEG analysis of ACT LC11R against untreated LC11R cells were mentioned on the top of the columns. Downregulated genes were marked in green, and upregulated genes were marked in red. For the autologous POC-ACT 1:1-E: T ratios were used of expanded and stimulated PBTs and TITs, as shown in the experimental setting, in **Figure 3D**. The ACT was performed for 4 days, whereby the cytokine profiles **Figure 3E** were determined on the day of ACT and flow cytometry, shown in **Figures 3K, L M and N** was determined for the different cell types at day 1 and day 4. **Figure 3F** shows cell viability over the ACT period. Figure G shows representative captures of 2D cultures of LC11R cells after 4 days of ACT, stained for the indicated molecules. Statistical outputs are p-values were plotted in the graph as follows: * p < 0.05; ** p ≤ 0.01 and *** p ≤ 0.001, **** p ≤ 0.0001.

### Enriched effector cell subsets derived from patient material fail to induce sufficient cell lysis in parental margin immune suppressive, proliferative TME

Next, patient-derived PBTs and TITs were employed for ACT, as shown in **Figure 3D**. Before ACT, immune cells were cultivated for 3 days with 180 IU/ml IL-2 and CD3/CD28 beads for stimulation. On day 3, cytokine and chemokine levels were determined, as shown in **Figure 3E**. Elevated levels of pro-inflammatory cytokines were detectable in both fractions. In reaction to endogenous IL-2 administration, PBTs synthesized elevated quantities of IL-17A, IL-6, IL-10, and IP-10 compared to the tumor tissue-associated counterparts.

Conversely, TITs generated elevated levels of free TGF-beta, IL-12p70, and IL-4. Nonetheless, both compartments do not consistently affect the LC11R cells, as evidenced in **Figure 3F** by observing cell proliferation following T-cell application. During the initial two days of treatment, both fractions demonstrated the capacity to inhibit proliferation to a certain degree; however, this impact was entirely nullified by day three. During the brief period of the suppressive impact, TITs demonstrated greater efficacy, consistent with the cytokine profile assessed before ACT. Contrary to prior findings, as depicted in **Figure 3G**, representative images of the co-culture with patient PBT 48 hours post-transfer demonstrate that LC11R cells efficiently inhibit activated immune cells by activating PDL-1, among other factors. Immune cell fractions extracted on day 0 and day 4 elucidate the T-cell subsets implicated in the immune-related effect, as illustrated in **Figure 3K–N**, where TITs are depicted in **panels K and L**, and PBT are presented in **panels M and N.** The cytotoxic CD8 T-cell subsets were categorized into activated, circulating, suppressive, and non-suppressive subpopulations. **Figure 3K** illustrates that TITs were recently active even after 4 days of co-culture. Unexpectedly, a more significant proportion of TITs exhibiting non-suppressive characteristics compared to the baseline timepoint was observed. This was likewise applicable to the circulating TITs. Upon elucidating the CD4 T-cell subsets, a similar picture was observed, wherein nearly all CD4 T-cells were recently activated, and the proportion of circulating CD4 T-cells climbed to 60%. At the same time, the suppressive CD4 T-cells were almost absent. Classifying the PBT determines that CD8 T-cells predominantly comprised T effector memory cells, consistent with the initial fraction. CD4 T-cells exhibited similar characteristics; however, CD8 T-cells demonstrated an even more significant proportion of TEM cells 4 days post-co-culture. TEMRA cells were not detectable, as in the initial fraction. The categorization of T-cell subsets and cytokine expression did not align with the T-cell-mediated effect observed for the LC11R co-culture of the parental margin tumor cells. However, patient-derived enriched and stimulated immune cells did not sufficiently eradicate the highly suppressive LC11R cells, suggesting that the front region exhibits a very specific adaptation and evasion strategy in the presence of autologous immune cells.

### Accessing the parental center TME by DEG-based pathway analysis reveals a low proliferating, lower inflammatory response and the inhibition of immune cell migration

In the second part of the study, we elucidate if the conducted specific adaptation could work vice versa and therefore enforces the autologous tumor-derived immune cells to eradicate the center region derived cells. Therefore, the tumor center tissue-derived cells were used for orthotopic transplantation to generate the tumor center cell line LC11Z. Next, a comprehensive analysis was conducted between the tumor front or margin and the center region to enhance the classification of the two regions, as described in **Figure 4**. **Figure 4A** illustrates a distinct pattern of diminished transcriptional activity at the center of the patient’s tumor. Substantial downregulation was observed for HGF, TRAM1, CREB4, and MPDZ, which govern proliferation and cell adhesion. Expression levels of genes associated with extracellular matrix formation and signaling and immunological and inflammatory responses such as IL1RAP and PTPN11 were diminished. Next, a comparative pathway analysis utilizing DEGs was conducted to compare the distinct regions against healthy hepatocytes. The objective was to classify regional disease-related disparities, particularly within the canonical pathways. The comparative results are displayed in **Figure 4B**. Here, canonical pathways associated with HCC progression and immune responses were analyzed. The disease-related pathway activation in both regions was nearly equivalent. Notably, IL-27 signaling was predicted to be activated in the center and inactivated in the margin. The activation of IL-27 signaling was shown to reduce the ability of immune cells to migrate and was linked to the inhibition of IL-2 production [29, 30].

**Figure 4:**
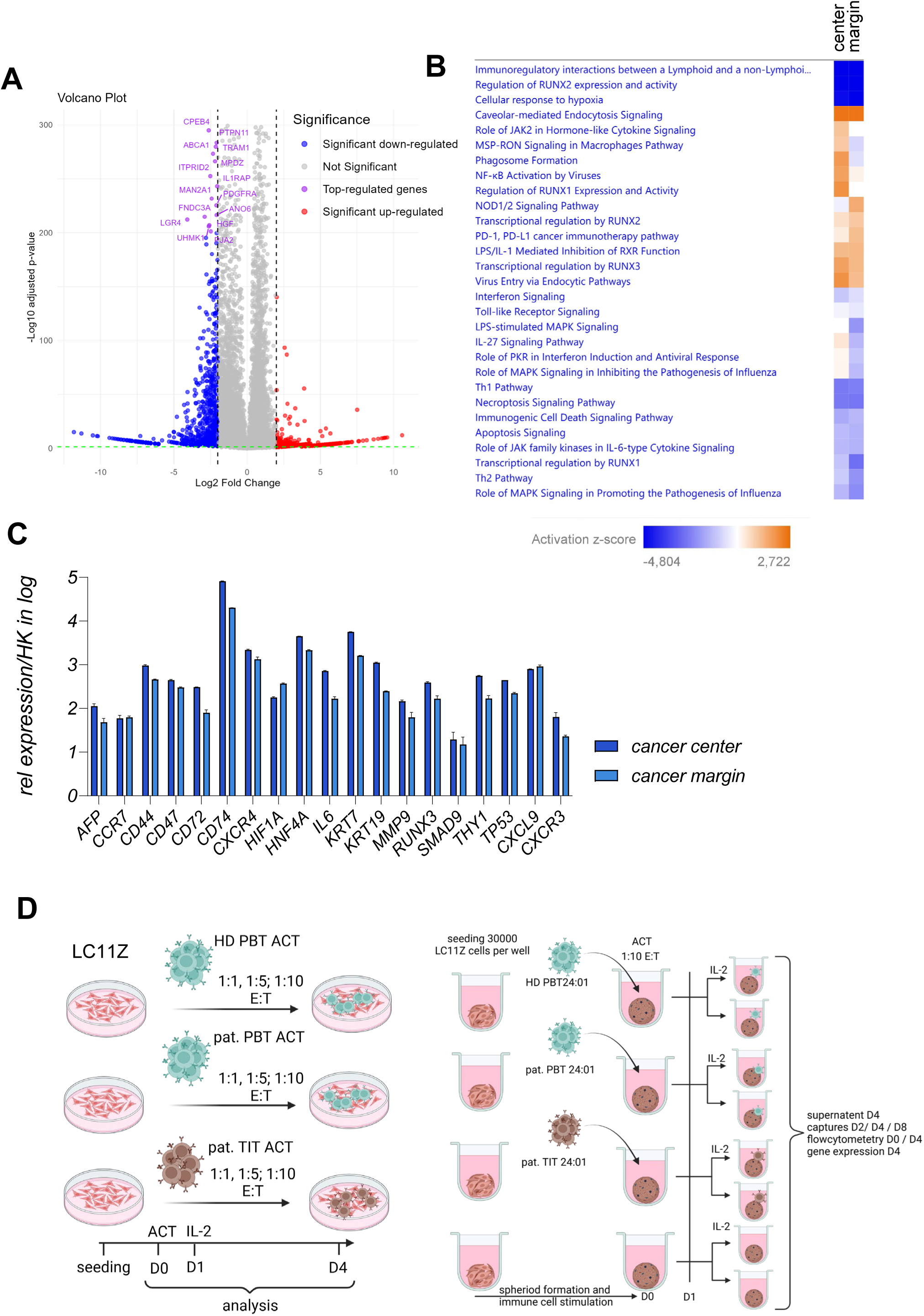
Comparative DEG-based pathway analysis of margin and center material. In **Figure 4A**, the volcano plot represents the DEG between the center and the margin region of the parental tumor used in the study, with indicated significances. The top-regulator genes are plotted in purple and marked with ensemble names. In **Figure 4B**, DEG-based comparative pathway analysis is shown by activation prediction. For comparative analysis, the margin and the center RNAseq analysis was compared to healthy hepatocytes (PHH) for normalization. The normalized data sets were used to perform the comparative canonical pathway analysis. In **Figure 4B**, the heatmap represents the indicated range of z-scores, whereby blue represents the prediction to be inactivated and orange to be activated. **Figure 4B**, the highest and lowest z-scores, with the indicated pathway names, were filtered and present. **Figure 4C** shows a more detailed snapshot of the DEG count data for the mentioned specific genes normalized against RPL0. **Figure 4D** shows the experimental settings for the conventional 2D and 3D cultures. As pathway analysis reveals the IL-27 signaling predicted to be upregulated in the center area, the IL-2 re-stimulation was included into the experimental setting as well as the different E:T ratios, since PDL-1 was predicted to be less active in the center and NOD2 signaling was predicted to be downregulated. Seeding and treatment were performed as shown, the supernatant was collected, and flow cytometry was performed at the indicated time points. Statistical outputs are p-values were plotted in the graph as follows: * p < 0.05; ** p ≤ 0.01 and *** p ≤ 0.001, **** p ≤ 0.0001.

Interestingly, the activation of the PD-1/PD-L1 pathway is predicted with a higher positive z-score in the margin. Therefore, the center is expected to be less inhibitory active [31]. This finding was interesting since the suppression of patient-derived immune cells was linked to higher PDL-1 expression in the margin cell line, shown in **Figure 3G**. RUNX3 exhibit a heightened transcriptional regulation in the center region. RUNX3 is a multivariable transcription factor that impacts T-cell differentiation but also the migration of tumor cell by induction of CD44 [32]. Interferon signaling was predicted to be inactivated in both regions, with a more significant inactivation observed in the center. The regulation of RUNX1 transcription and activity was very pronounced in the center but undetectable in the margin. The NOD1/2 signaling pathways exhibited distinct activity scores, demonstrating heightened activity at the margins and suppression at the center, as shown in **Supplementary Figures 4 and 5**. NOD2 is activated by viral RNA recognition and leads to enhanced production of inhibitory cytokines, such as IL-8 [33]. A single gene pattern analysis, illustrated in **Figure 4C**, indicates that most specified genes had reduced expression in the margin, except for CCR7, HIF1alpha, and CXCL9. Both regions expressed high levels of CD74, CK7, CK19, HNF4A, and CXCR4. CD44 gene expression was determined in both areas; however, as seen in **Supplementary Figure 1**, there was no surface presentation of CD44 on LC11R cells. Nonetheless, parental tumor center cells were utilized as previously described to generate LC11Z cells, as depicted in **Figure 1A**. LC11Z cells were utilized for ACT, as illustrated in **Figure 4D**.

### Patent-derived PBTs sufficiently suppress proliferation of tumor center derived LC11Z cells at low E:T ratios and additional IL-2 application

To examine the T-cell response and the cytolytic capabilities of immune cell fractions obtained from the patient, patient-derived center cells were used for ACT, as illustrated in **Figure 4D**. Initially, HD24:02 PBTs were used to confirm our findings. **Figure 5A** illustrates that, in the presence of tumor center cells (LC11Z), HD24:02 PBTs were able to inhibit proliferation within the initial 24 hours when utilizing elevated effector to target (E:T) ratios.

**Figure 5:**
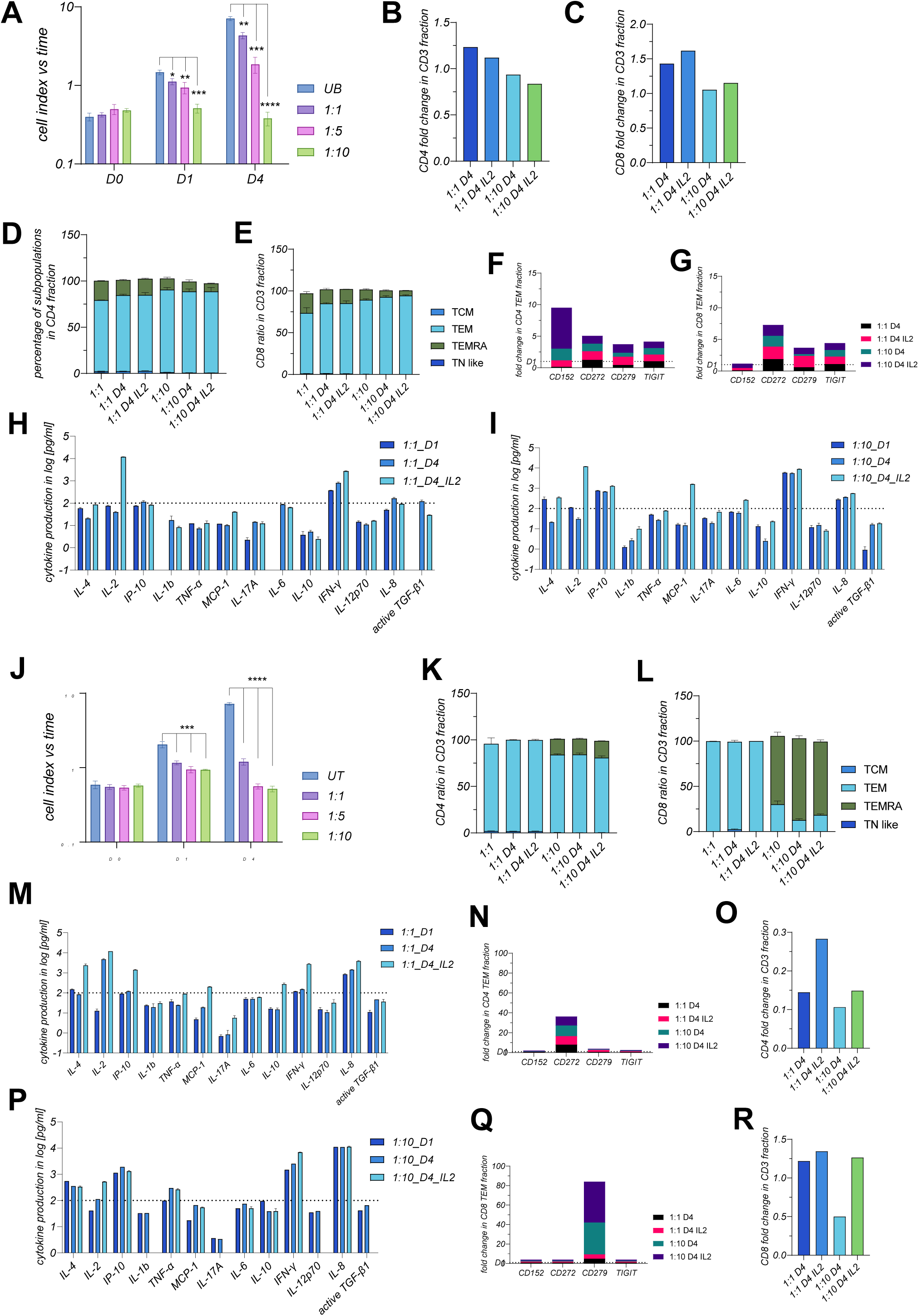
Adoptive allogenic and autologous PBT transfer using cancer center-derived cells. For allogenic ACT, shown in **Figures 5A – I**, PBTs were stimulated for 3 days and transferred in indicated ratios. After 24 hours of co-culture, 180IU/ml of IL-2 were additionally applied in the re-stimulation groups. In **Figure 5A**, cell viability was measured and is plotted for the groups receiving additional IL-2. **Figures 5B – I** show both re-stimulated and stimulated groups in comparing low and high E:T ratios. In **Figures 5B – G**, flow cytometry-based characteristics are plotted in fold change to parental or were determined in percentages of the indicated populations. T cell differentiation was classified in the CD3+CD4+ or the CD3+CD8+ subpopulation by applying CCR7 and CD45RA for differentiation of T-naïve, T-central, T-effector memory and T-effector memory representing CD45RA. The present inhibitory marker is plotted in fold change to the initial PBT signature after 3 days of stimulation. For cytokine and chemokine signatures, low (**Fig. 5H**) and high (**I**) E: T PBT-ACTs are presented at different time points and for different stimulation levels, as indicated. In panels **Figures 5J – R**, the autologous PBT ACT is plotted in direct comparison to **panels A - I**. Statistical outputs are p-values were plotted in the graph as follows: * p < 0.05; ** p ≤ 0.01 and *** p ≤ 0.001, **** p ≤ 0.0001.

The re-stimulation by additional administration of endogenous IL-2, 24 hours after ACT, failed to facilitate cytolytic activity in 1:1 or 1:5 ratios, similar to the results observed with the margin-derived LC11R cells, as illustrated in **Figure 2B**. T-cell differentiation was investigated in CD8 and CD4 subsets at 24 and 96 hours after T-cell transfer for 1:1 and 1:10 E:T ratios, with or without re-stimulation at day 1. **Figures 5B and C** illustrate that CD4 T-cell growth is inhibited by the administration of IL-2 and increased quantities of T-cells.

Conversely, the proliferation of CD8 T-cells was augmented by IL-2 administration; however, like CD4 T-cells, proliferation was diminished by applying elevated quantities of effector cells. CD4 T-cells mainly differentiate into TEM and partially into TEMRA cells 24 hours post-transfer, exhibiting a reduced fraction of TEMRA cells in 1:10 ratios. The additional administration of IL-2 does not alter the differentiation profile of CD4 T-cells. On day 4 post-transfer, the TEMRA subset diminished in both E:T ratio conditions. The identical results have been observed for the CD8 T-cell subset. TEMRA cells were reduced in increasing quantities, with the most significant reduction occurring in the 1:10 condition following the administration of IL-2 on day 1. **Figures 5F and G** demonstrate that the expression of the inhibitory marker was elicited in all situations relative to day 1 for CD4 and CD8 T-cells. CD272, TIGIT, and CD279 (PD-1) were significantly upregulated on CD8 T-cells, although CD152 exhibited no increase. All detected inhibitory molecules on CD4 T-cells were elevated, with CD152 being the most prominent in the 1:10 IL-2 conditions. An IL-2-associated rise pattern was observed in CD4 and CD8 T-cells for CD279 (PD-1) and CD152, with CD152 less represented on CD8 T-cells than the original assessment on day 1. The 10-fold T-cell application generally yields increased cytokine and chemokine production, as **Figures 5H and I** illustrate. The cytokine profile indicates higher IFN-γ production concurrent with increased IL-2 levels on day 4. Significant production of IP-10, IL-6, and IL-8 was determined, with or without IL-2. In the 1:10 T-cell application, elevated levels of IL-17A and IL-10 were observed compared to the 1:1 condition, highlighting the elevation of the inhibitory marker depicted in **Figure 5F**. IL-1beta was exclusively observed in the 1:1 condition on day 4, whereas in the 1:10 condition, IL-1beta was discovered in negligible quantities on day 1 and increased by day 4. IL-4 was generated in an IL-2-dependent manner under the specified conditions. Nonetheless, LC11Z cells effectively inhibit T effector cells in reduced quantities while achieving enough immune evasion.

Subsequently, we examine if the immunological profile of T-cells obtained from the patient’s blood may counteract immune suppression by creating a stimulatory environment. Consequently, the patient’s PBTs were administered in 1:1, 1:5, and 1:10 ratios under identical settings for the same time intervals and examined the T-cell fractions. Here, patient PBTs effectively suppressed proliferation until day 1 and, following IL-2 administration, exerted an enhanced cytolytic impact, as illustrated in **Figure 5J**. **Figure 5K** illustrates that the CD4 T-cell fraction predominantly consisted of TEM cells at lower quantities. However, a ten-fold increase in T-cell numbers resulted in a transition from TEM to TEMRA differentiation. Like the CD8 fraction, as illustrated in **Figure 5L**, where TEMRA cells represent the most significant differential condition within the CD8 T-cell fraction. The increased prevalence of TEMRA cells in the 1:10 fraction likely enhanced the cytolytic effect observed in **Figure 5J**. Upon assessing the inhibition profile of the patient’s T-cells, a notably consistent increase of CD272 expression in the CD4 T-cell subset was observed, although other inhibitory markers were expressed at equivalent levels or lower compared to day 1, as illustrated in **Figure 5N**. On CD8 T-cells, CD279 (PD-1) was markedly elevated, as illustrated in **Figure 5Q**. Notably, the 1:10 condition, predominantly exhibited by TEMRA cells, induces the highest expression of CD279 (PD-1). The cytokine profiles illustrated in **Figures 5M, and P** demonstrate a distinct pattern in the healthy donor T-cells, producing elevated levels of IL-8 rather than IFN-γ. Reduced IL-17A levels and increased IL-10 production were determined. IL-4 was produced in elevated quantities, and notably, the T-cells of patients generated substantial levels of IL-2. As illustrated in **Figures 5O and R**, CD4 T-cells exhibited a reduction on day 1, whereas CD8 T-cells demonstrated proliferation on day 1 and continued to do so following IL-2 administration until day 4. This is notable despite the predominant T-cell type’s brief lifespan, which accounts for the 50% decrease by day 4 in the absence of IL-2 treatment. The drop of CD4 T-cells cannot be attributed to the rapid demise of TEMRA cells, a decline was observed in fractions where TEMRA cells were absent. We propose a significant suppression of proliferation attributed to CD272. A distinct suppression pattern was seen in the patient’s T-cells, which was lacking in healthy donor T-cells, indicating that the immunological signature of the patient’s immune cells facilitates a highly conserved evasion of tumor cells.

**Figure 6A** illustrates gene expression analysis conducted after 4 days of co-culture with depicted PBTs. The CD4 expression level was elevated in patients’ T-cells. However, CD8 expression was markedly low. Notwithstanding the cytokine and chemokine levels illustrated in **Figure 5**, the gene expression of IFN-γ was most pronounced in the specimens containing patient PBTs. The migratory marker CCR7 exhibited elevated expression levels in the patient-derived PBTs, while CD44 demonstrated a reduced expression pattern. Nonetheless, these findings corresponded with the differentiation status of the T-cell subsets. A reduced level of HNF4A, prominently expressed in tumor center cells, as shown in **Figure 4C**, indicates the tumor cell lysis, supported by Caspase 1 and 3 inductions. HNF4A expression was diminished when utilizing T-cells from healthy donors, which also exhibited a more pronounced induction of Caspases 3 and 1, indicating that T-cells from healthy donors and patients could induce apoptosis and inflammation, thereby diminishing tumor cells originating from the center.

**Figure 6.**
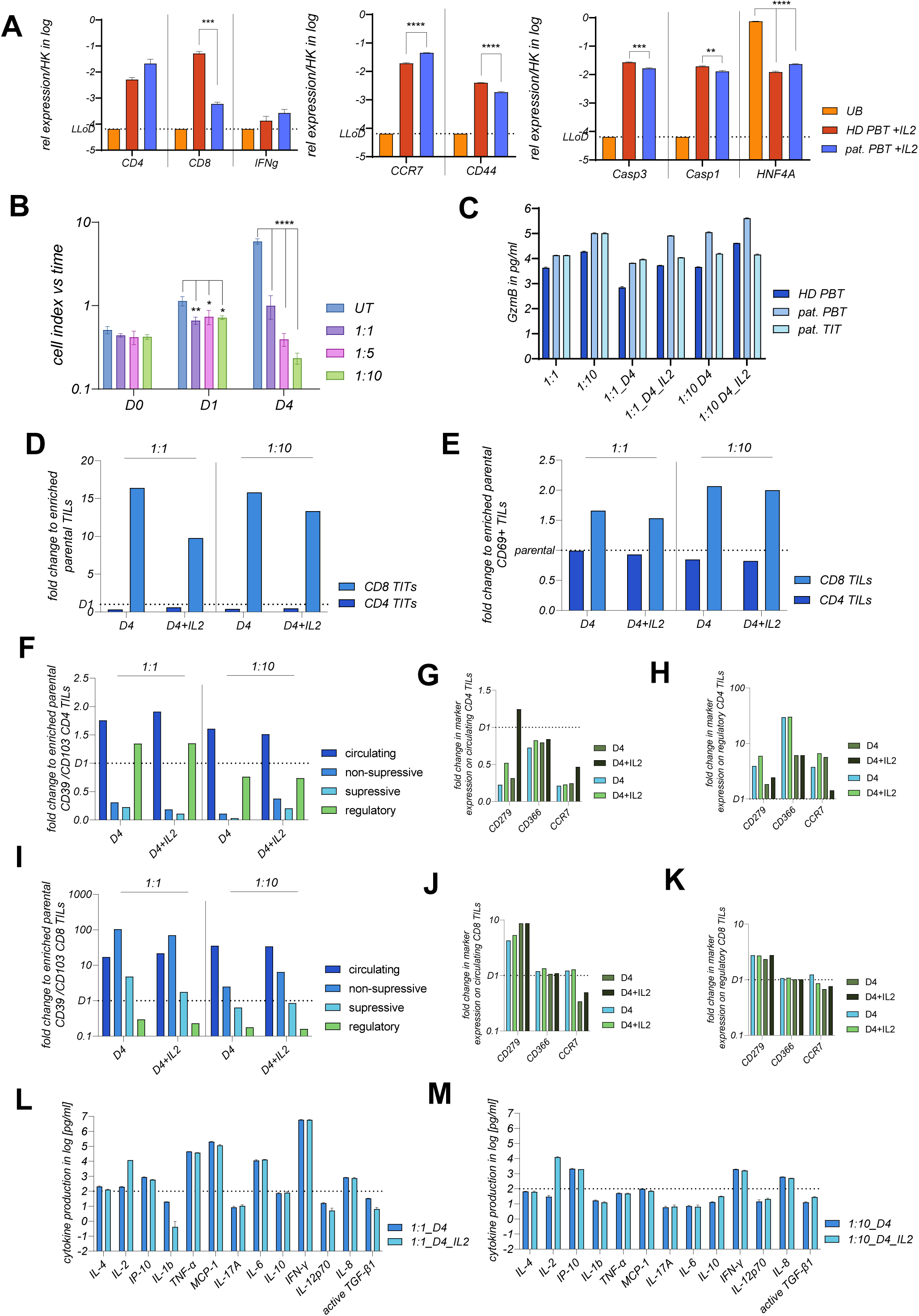
PBT-ACT induced gene expression pattern and the impact of the autologous TIT-ACT on center cancer cells. **Figure 6A** shows the gene expression patterns for the PBT-ACT, for the indicated genes and groups. Untreated LC11Z cells or allogenic and autologous PBT-ACT co-cultures undergoing re-stimulation were isolated and employed for TaqMan-based amplification of T cells or cancer-specific genes. For each isolation 4 biological replicates were pooled to reach a sufficient number of cells. Expression was normalized against housekeeper gene RPL0 and plotted in a logarithmic scale. **Figure 6B**, represents the autologous TIT-ACT with indicated E:T ratios, undergoing the experimental settings illustrated in **Figure 4D**. In **Figure 6B**, longitudinal cell viability acquisition after TIT-ACT is presented. Re-stimulation was performed on day 1 after ACT for all ratios, as described before. **Figure 6C** shows the GZMB production of the PBTs- and TITs during ACT, in the supernatant over time and conditions is displayed in pg/ml. For the panels, **Figure 6D – K**, flow cytometry was performed on day 0, 1 and 4 for the stimulated and re-stimulated TITs. In **Figure 6D**, the fold change analysis for the CD3+ subpopulations are plotted, while in **Figure 6E**, the recently activated (CD69+) subpopulations are plotted in fold change to the initial TIT-fraction. In **Figures 6F and I**, either CD4 **(F)** or CD8 **(I)** TIT subpopulations are illustrated over time and conditions. The inhibitory signature for CD366 and CD279 (PDL-1), as well as the migratory marker CCR7, is presented for the indicated TIT differentiation types (circulating (**Figures 6G and J**) and regulatory (**Figures 6H and K**)) in fold change for CD4 TITs in the **Figures 6 G and H** and for CD8 TITs in the **Figures 6 J and K**. The cytokine and chemokine production were measured and plotted in **L** 1:1 and in **Figure 6M** for 1:10 conditions in stimulation (D4) and re-stimulation (D4_IL2) of TITs used for the ACT in LC11Z cells. Statistical outputs are p-values were plotted in the graph as follows: * p < 0.05; ** p ≤ 0.01 and *** p ≤ 0.001, **** p ≤ 0.0001.

### TIL-derived CD8 T-cells exert cytolytic and suppressive functionality in their familiar TME

Next, we utilized enriched and activated TITs for T-cell transfer, as illustrated in **Figure 4D**.**Figure 6B** demonstrates that varying T: E ratios showed no differences in inhibiting tumor cell proliferation on day 1, even when accomplished by higher GZMB levels, as shown in **Figure 6C**. Suppression was more successful than with PBTs from HD24:02 and equivalent to that achieved with the T-cell fraction obtained from the patient’s blood, which was in line with the GZMB titer. Following the administration of IL-2, TITs demonstrated the capacity to inhibit proliferation and promote cell death at an equivalent or even superior level compared to PBTs, as illustrated in **Figure 5**. Interestingly, GZMB production was most prominent and constantly increased when treating LC11Z cells with patient’s PBTs, while TITs produced a consistent but less increased amount of GZMB. **Figure 6D** illustrates that the percentage of CD8 TITs increased across all settings; however, the increase was notably less pronounced with the application of IL-2. As illustrated in **Figure 6E**, the CD3+ CD8/CD69 expressing cells increased by up to 2-fold over time. Still, the CD4/CD69 expressing cells diminished in comparison to the parental TITs utilized in the cell co-culture. The application of IL-2 does not augment but somewhat reduces the proportion of the CD69 positive fraction. **Figure 6F** illustrates that regulatory TITs were diminished by an elevated T: E ratio. We observed a distinct downregulation in the subset of non-suppressing and suppressing cells following treatment.

In summary, the most significant downregulation of regulatory, suppressive, and non-suppressive CD4 TITs was determined in the 1:10 condition. A distinct trend was visible for CD8 TITs after 4 days of therapy, as illustrated in **Figure 6I**. As shown, we obtained a significant rise in the percentage of CD8 TITs in circulating cells, accompanied by a substantial drop in regulatory cells across all conditions. Overall, non-suppressing CD8 TITs were elevated on day 4 relative to day 1, particularly at a 1:1 ratio. Reduced expression of CD279 (PD-1) and CD366 was determined on circulating CD4 TITs, as illustrated in **Figure 6G**, along with a reduction in CCR7 within this population. In regulatory CD4 TITs, as shown in **Figure 6H**, upregulation of CD279 (PD-1), CD366, and CCR7 was detectable. The induction of CD366 was independent of IL-2 administration in the regulatory fraction, although correlated with circulating CD4 TITs. The inhibitory marker CD279 (PD-1) was elevated on both CD4 and CD8 TITs across all conditions, but CD366 was expressed at levels comparable to those acquired on day 1, as displayed in **Figures 6G-K**. A steady expression of CCR7 was detected relative to day 1 in the 1:1 E:T settings, as illustrated in **Figure 6K**, whereas CCR7 expression diminished in the 1:10 E:T conditions. On day 1, no non-suppressive CD8 TITs were observed; however, following 4 days of treatment, about 10% of all CD8 TITs were identified as non-suppressive (data not shown). This subpopulation had elevated CD366 and CD279 (PD-1) expression, observed in 30-40% of the subset, as illustrated in **Supplementary Figure 6**. Simultaneously, the non-suppressive CD8 TITs exhibited CCR7 expression in 60-70% of cases.

The cytokine and chemokine profiles of the various conditions depicted in **Figures 6L, and M** indicate that the production of IFN-γ, TNF-alpha, MCP-1, IL-4, and IL-6 was more pronounced in samples with reduced TIT quantities, whereas IP-10 levels were elevated when utilizing tenfold more TITs compared to LC11Z cells. Notably, IL-8, which was visible in elevated concentrations during the PBT studies, exhibited comparatively low levels in the measured amounts. Collectively, the same differentiation profile was detected for CD8 TITs irrespective of the target T-cell line, the quantity of effector cells, or the re-stimulation and after four days; however, there was a heightened production of IL-1beta, free TGF beta, and IL-2 when the margin cell line served as the target T-cell. Notably, similar to PBT, the suppression of CD8 positive cells was achieved by the expression of CD279 (PD-1), unlike allogenic T-cells. Nonetheless, the utilization of TITs and PBTs sourced from the original tumor and blood proved more effective in lysing tumor cells or, at a minimum, inhibiting growth in a 1:1 scenario, as illustrated in **Figure 7A**.

**Figure 7:**
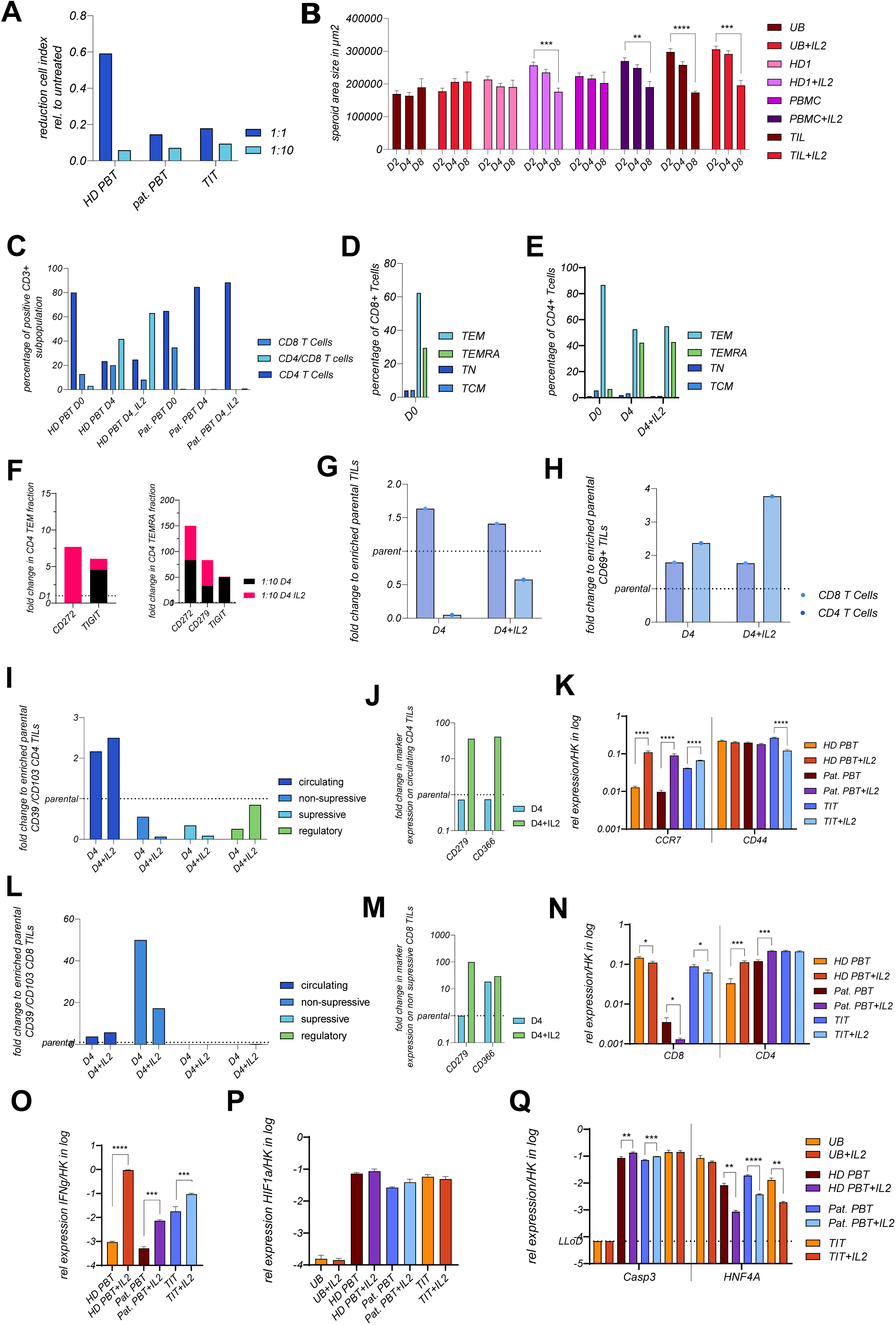
Comparative modelling of allogenic and autologous PBT- or TIT-ACT in 3-dimensional tumor microenvironment using protein and gene expression analysis. For the assessment of migratory barriers, the formation of LC11Z spheroids was established before ACT was performed. As described before, the most sufficient ACT E: T ratios using LC11Z cells were observed in 1:10, as summarized by calculating the fold change shown in **Figure 7A**. Therefore 1:10 ratios were used in the 3D ACT using PBTs and TITs, as illustrated in the **Figure 4D**, right panel. IL-2 re-stimulation was performed on day 1 after ACT. In **Figure 7B**, the automatically calculated spheroid areas are plotted over time and depend on the conditions and the indicated groups. The ACT followed until day 8 for the area calculation of the spheroids, also shown in the supplementary Figure 7 and described in the method section. For flow cytometry, immune cells that had not migrated into the spheroids were collected on day 4 after ACT and analyzed as indicated. In C, the allogenic and autologous PBTs were analyzed on day 4 after ACT and were plotted in the percentage of CD3 T cells in PBTs and TITs compared to the initial immune cell fractions. In **Figures 7D and E**, the autologous CD4 and CD8 PBTs were analyzed on differentiation level over time. Since CD8 PBTs were not detectable in the CD3 fraction, only the initial CD8 fraction is plotted in **Figure 7D**. In **Figure 7F**, the inhibitory signature of autologous PB CD4 TEM (left) and TEMRA (right) is plotted for a dateable marker in fold change to the initial presentation on day 0. Panel G presents the fold change in CD4 and CD8 TITs after ACT compared to the initial parental immune cell fraction used for TIT-ACT, while in **Figure 7H**, the recently activated TIT subtypes are plotted in fold change, as described before. CD8 and CD4 TIT subpopulations were analyzed for their differentiation status shown in the panel I and L. Cell types which were most prominent were visualized for the expression of CD366 and CD279 (PDL-1) for CD4 TIT in J and CD8 TIT in M. All comparisons were made using the parental stimulated immune cell fraction at day 0 or the immune cell fraction on day 1 after ACT, as visualized by the dotted line in the graphs and indicated on the y-axis. For the specific gene expression pattern, 3 spheroids of each condition and cell type were collected and used for the 2-step TaqMan PCR analysis, for gene expression level determination of the indicated genes, shown in panel K, N and O – Q. As described before, relative expression was calculated using the housekeeper gene RPL0. Statistical outputs are p-values were plotted in the graph as follows: * p < 0.05; ** p ≤ 0.01 and *** p ≤ 0.001, **** p ≤ 0.0001.

### TIL-derived CD8 T-cells effectively migrated into 3-dimensional tumor center spheroids and achieved cytolytic functionality, while PBMC derived CD8 T-cells diminished even with IL-2 application

Subsequently, to examine the migratory potential of the three cell types, we conducted transfer experiments utilizing spheroids derived from LC11Z cells. Spheroids were that undergo a three-week development period, applying identical parameters as previously published, utilizing 1:10 ratios. Immune cells and spheroids were harvested and examined using flow cytometry and specific PCR on day 4 post-transfer. Spheroid regions were captured, and area size was calculated automatically on days 2, 4, and 8 post-transfer. Representative captures are illustrated in **Supplementary Figure 7**. **Figure 7B** illustrates that spheroids treated with IL-2 or utilized for co-culture demonstrate increased area sizes on day 2, with further enhancement shown after re-stimulation of immune cells, indicating the migration of immune cell fractions into the spheroids. This observation aligns with the more significant percentage of CCR7-positive immune cells in the 1:10 ratios. Following four days of co-culture, a slight decrease in spheroid size was determined when LC11Z cells were treated with T-cells generated from healthy donors or patients’ PBTs without re-stimulation.

In contrast, spheroids treated with TIT exhibit a more significant reduction. However, using IL-2 results in enhanced migration and eventual cytotoxicity, as evidenced by a decrease in size observed on day 8 post-treatment. On day 4, the T-cell subset was delineated, as illustrated in **Figure 7C**, by diminished CD8 and CD4 levels, with the CD8 fraction further reduced upon applying IL-2. This discovery aligns with the augmented migratory potential of immune cells. Conversely, healthy donor T-cells exhibited a significant proportion (40-70%) of CD4/CD8 positive T-cells. This characteristic was absent in patient-derived T-cells, as illustrated in **Figure 7C**. The patient-derived CD8 T-cells initially comprised TEMRA and TEM cells, as seen in **Figure 7D**, which were undetectable in the spheroid supernatant after 4 days. Detectable CD4 T-cells exhibited a higher number of TEMRA cells than the original immune cell fraction, with TEM comprising nearly the entirety of the CD4 fraction. The fold change expression of the inhibitory marker, depicted for CD4 TEM and TEMRA cells in **Figure 7F**, demonstrates the induction of CD272 on CD4 T-cells with and without the additional application of IL-2. Still, TIGIT was only elevated when IL-2 was administered to the co-culture. In CD4 TEMRA cells, CD272 was significantly upregulated, along with CD279 (PD-1) and TIGIT. TIGIT was assessed solely on unstimulated CD4 TEMRA cells. On day 4, the gene expression levels of CD4 and CD8 from isolated spheroids indicate that CD8 expression was minimally detectable when transferring patient-derived T-cells, as illustrated in **Figure 7N**, whereas CD4 expression was observed at elevated levels relative to healthy donor co-culture samples.

**Figure 7G** illustrates that the patients’ TIT fractions exhibited a significant decrease in the TIL-derived CD8 T-cell fraction, whereas the CD4 T-cells increased relative to the starting cell compartment. In contrast to T-cells from the healthy donor, and consistent with the patients’ T-cells, 2% of the CD3 positive fraction were characterized as CD4/CD8 TITs (data not shown). Consequently, the elevated quantity of CD4/CD8 double-positive cells was exclusively noted in the healthy donor’s PBTs in 3D conditions. **Figure 7H** illustrates that recently activated TITs increased until day 4 compared to the initial compartment, with activated CD8 TITs being particularly pronounced when re-stimulated. Further analysis of the differentiation status of CD4 and CD8 TITs was conducted, which aligns with the composition of CD4 TITs in 2D culture under 1:10 conditions, as illustrated in **Figure 7I**. Circulating CD8 TITs were present in lower quantities compared to the 2D culture, indicating their migration inside the spheroids. Non-suppressive CD8 TITs significantly increase the proportion, as illustrated in **Figure 7L**. Circulating CD4 TITs have diminished levels of the inhibitory markers CD279 (PD-1) and CD366 in the absence of IL-2 but show an up to 40-fold increase with IL-2 administration. In the most prominent CD8 TIT subset, the non-suppressive cell, IL-2 administration elevated the expression of CD279 (PD-1) and CD366, whereas cells not re-stimulated did not exhibit increased expression of CD279 (PD-1).

The proposal of actively migrating immune cells into the spheroids was emphasized by the gene expression of CCR7 depicted in **Figure 6K**, which indicated a greater expression rate with IL-2 supplementation. For TITs elevated CCR7 expression was detected in comparison to PBT-ACT, even in the absence of IL-2 supplementation. No changes in the expression of the migration marker CD44 could be detected except for the downregulation when applying TITs and IL-2. To analyze the migratory capacity, gene expression related to CD4 and CD8 was conducted, and reveals the maximum level of CD4 expression, irrespective of IL-2, while CD8 expression diminishes under IL-2 conditions. Patients’ CD4 PBT-cells exhibited a much higher migration rate into the LC11Z dense formations compared to CD8 PBT-cells, which were either undetectable or minimally detectable. Tissue-derived T lymphocytes exhibited the ability to migrate into the spheroids with elevated expression levels of CD4 and CD8, as illustrated in **Figure 7N**. The detection of IFN-γ expression under various settings, shows that IL-2 treatments were effective, as described in **Figure 7O**. The expression level of HIF1alpha, linked to a hypoxic environment, significantly increased independent of the additional IL-2 administration following immune cell transfer, as illustrated in **Figure 7P**. Elevated levels of caspase 3 were determined accompanied by HNF4A reducion. Caspase 3 expression appeared to be significantly increased for PBTs when applying IL-2, while TIT ACT results in IL-2-independent high expression levels of Caspase 3. However, the decrease in the hepatocyte marker HNF4A was not directly associated with the expression of caspase 3. The findings related to the spheroid structure signify the initiation of apoptosis. Consequently, T-cells from the HBV-HCC patient were capable of causing cytolysis through IFN-γ, GZMB, and the death receptor ligand, thereby triggering ferroptosis, proptosis and apoptosis, with TIT being even more effective in lower cell numbers or without IL-2 re-stimulation. Under these conditions, CD4 TITs exhibit lower inhibitory marker expression, CD8 TITs were expanded during ACT and producing high IFN-γ and constant GZMB levels.

## Discussion

The quest for effective adoptive cell transfer strategies in treating HCC, particularly in HBV-related cases, necessitates a detailed examination of T-cell sources and their functional characteristics within a highly suppressive TME in the individual patient [34, 35]. Understanding individual barriers and opportunities is paramount since, despite interventions, the recurrence rate within a five-year period remains alarmingly high, reaching up to 70% [36]. Despite extensive efforts immune checkpoint blockade therapies, therapeutic efficacy remains suboptimal, often hindered by treatment resistance, with an overall response rate to these interventions below 30%. In sum, the clinical need for further advancements remains significant in uncovering the mechanisms of the individual tumor microenvironment in the presence of cell-based immune therapies. Our study focuses on these distinct behaviors of an individual TME, in the presents of transferred CD3 T-cells derived from the patient-derived tumor-infiltrating lymphocytes (TIT) and peripheral blood mononuclear cells (PBT) in comparison to T-cells derived from peripheral blood mononuclear cells of an HLA-matched healthy donor. We could show that TITs provide a specific source to overcome the TME migration barrier for cell-based immunotherapy in an individual HCC stetting. Moreover, our findings advised on implementing an individual evaluation of resected tissues to predict the efficacy of immune therapies in an individual setting, in a likely scenario, the reoccurrence.

TIL therapy is based on the activation and expansion of infiltrating T-cells extracted from the tumor itself (TITs), which are intrinsically reactive against tumor antigens, setting an exciting rationale for its use against the changing and heterogeneous cell population of solid tumors [16]. However one of the biggest challenges for TIL-based ACT is the availability of the cell source itself which can be expanded [37]. In this study we examined the isolation and expansion process of TITs from the tumor material of 6 HCC patients. We were able to isolate adequate TIL cell numbers, which could be used for expansion. In line with others, our results showed that expanded and stimulated TITs exhibit the capacity to migrate into a highly suppressive TME, and exhibit anti-tumor activity, characterized by increased numbers of CD8 T-cells [14, 16, 17]. We showcased that the CD8 T-cell fraction predominantly consisted of non-suppressive and circulating T-cell subtypes, underscoring their potential efficacy in targeting tumor center cells.

Moreover, the upregulation of CD279 (PD-1) among these cells suggests an activated state, although it also raises concerns about the potential for exhaustion in the face of persistent tumor antigens [31]. The production of pro-inflammatory cytokines in high titers, including TNF-α, MCP-1, IL-6, and IFN-γ when using TITs, reinforces their role in driving robust immune responses. However, the study also showcased that autologous TITs and PBTs exhibited reduced CD8 T-cell activity in the tumor margin, pointing to a particular immunosuppressive tumor microenvironment. In the comparative analysis, allogenic PBTs demonstrated enhanced initial activation, as evidenced by the upregulation of IL-4, IL-2, IP-10, and IL-1β. While these cells showed effective responses at the tumor front region, their cytotoxic capacity in the center of the tumor was limited, e.g. by a lower Granzyme B production in direct comparison to the autologous PBTs, which indicates a lack of sustained anti-tumor efficacy. Furthermore, the healthy donor PBTs exhibited a decrease in both CD4 and CD8 T-cells after 3D co-culture, with an increase in CD8/CD4 double-positive cells. This indicates a transitional phase rather than a strengthened functionality, complicating their overall effectiveness against HCC.

When examining the T-cell behaviors under different culture conditions, stark differences were noted between 2D and 3D environments. In 3D cultures, autologous PBTs displayed a remarkable decrease in CD8 T-cells, while CD4 T-cells increased, hinting at a regulatory or compensatory mechanism in the face of TEM-like structure presence. Conversely, healthy donor’s PBT-cells demonstrated a similar downward trend in CD4 and CD8 populations, suggesting that the T-cell response becomes more heterogeneous and less effective in solid tumor contexts. The high and consistent induction of CD44 and HIF1α expression in treated spheroid cultures signifies a retained potential of T-cells to exhibit effective immune responses, as both markers support immune evasion and suppression in solid tumors [38, 39]. In line with these findings, we examined a sufficient migration of T-cells into the spheroids, executed by CD4 and CD8 expression patterns. The re-stimulation with IL-2 elevates CCR7 expression and enforces the migration capacity of transferred PBTs. Interestingly, TITs were able to express high CCR7 levels without re-stimulation.

In sum, the mechanistic adaptation of the TME towards migration and reduced oxygen availability (CCR7, CD44, HIF1α) points to the quick shift from proliferation to survival caused by the ACT. This milieu appears to be sufficient in the reanimation of the parental immune suppressive mechanism, leading to reduced cytolytic fractions in transferred PBTs. At the same time, TIT exhibit a stable resistance towards immune suppression.

Taken together, elucidating the nuances of T-cell behavior in a specific tumor microenvironment lays the groundwork for optimising individual ACT strategies for treating HCCs. Notably, autologous TITs demonstrate a solid capacity to produce pro-inflammatory cytokines such as TNF-α, MCP-1, IL-6, and IFN-γ, which are critical for mediating effective anti-tumor responses. In contrast, autologous PBTs exhibit immunological profiles that were less effective. However, both cell sources exhibit a less effective suppression of tumor cell proliferation in the tumor margin. This disparity highlights the challenges faced by endogenous immune cells within the immunosuppressive tumor microenvironment, particularly in areas where tumor cells exert the strongest inhibitory effects on T-cell function.

Interestingly, allogenic PBTs from healthy donors showed a remarkedly efficacy in the tumor margin, which raises important considerations about the nature of immune responses elicited by T-cells from different sources. The upregulation of IL-4, IL-2, IP-10, and IL-1β in allogenic PBTs facilitate enhanced T-cell activation and recruitment, contributing to their effectiveness in areas where general inflammatory conditions are necessary to combat immune evasion strategies. This response contrasts with the autologous TITs and PBTs, where the tumor microenvironment has dampened their cytotoxic functionality, evident in their inability to exert anti-tumor effects in the marginal area despite the presence of some pro-inflammatory cytokines. Moreover, the reduced production of Granzyme B in allogenic PBTs, suggests that effective cytotoxic capacity does not strictly correlate with cytokine production profiles. This indicates that healthy donor T-cells may adopt a more generalized activation status that does not fully translate into cytotoxicity against the specific HCC cells, likely due to varying sensitivities to the specific immunosuppressive signals in the tumor microenvironment. This suggestion was supported by the general upregulation of inhibitory markers determined for autologous PBT ACT.

Given that TILs are selected for their specificity and reactivity to specific tumor antigens during their development within the tumor microenvironment, their comparative struggle in the margin attests to the need for potent adjunctive strategies to overcome the tumor’s firewall. A focus on refining techniques to augment TIL activity, would be to evaluate combination therapies that leverage both autologous TILs and allogenic PBTs, to better harness the immune system’s potential to effectively combat HCCs, aiming for improved clinical outcomes in this challenging disease landscape.

## Author contributions

JK initiated and supervised the research study; JK, LS and MD designed the experiments; AH and KS supervised the collection of human samples; JK, LS, MCH, MV, MG and TV conducted experiments and acquired data; JK, LS analyzed data; JK, SL, SP, MD wrote the manuscript. All authors discussed the data and corrected the manuscript. All authors had access to the study data and had reviewed and approved the final manuscript.

## Acknowledgements.

We thank Tobias Gosau and Ursula Mueller for their excellent work, Claudia Dettmer, Eileen Maly, and Karina Boerner for their excellent technical assistance, and Kristoffer Riecken for providing the LeGO vector, which was used for stable transduction of LC11R_LUC cells.

## Aberration

ACT: adoptive cell transfer therapy
CD152: Cytotoxic T-lymphocyte-associated protein 4
CD272: B- and T-lymphocyte attenuator
CD279 (PD-1): Programmed cell death protein 1
HBV: Hepatitis B virus
HCC: Hepatocellular carcinoma
LC11R: liver cancer cell line 11 derived from parental margin tissue
LC11Z: liver cancer cell line 11 derived from parental center tissue
PBT: peripheral blood-derived T-cells
TCM: central memory T-cells
TEM: effector memory T-cells
TEMRA: Effector memory T-cells expressing CD45RA
TIGIT: T-cell immunoreceptor with Ig and ITIM domains
TIL: tumor-infiltrating lymphocytes-derived T-cells
TME: Tumor microenvironment
TME: tumor microenvironment
TN: naïve like T-cells

