## Supplementary Figure 1 for "Accessing the specific capacity of TIL-derived CD8 T-cells to suppress tumor recurrence in resectable HBV-HCC patients"

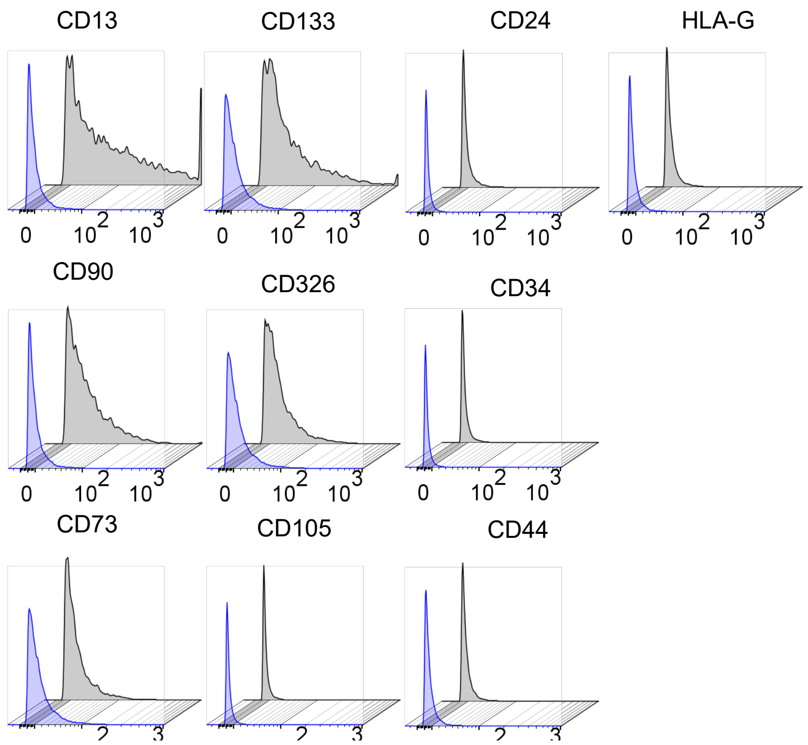

*Supplementary Figure Legends*

***Supplementary Figure 1:*** **Flow cytometry-based characterization of generated LC11R cells.**

Offset histograms represent the unstained LC11R cells in light blue and the stained LC11R cells in grey for the indicated molecules.
