## Supplementary Figure 2 for "Accessing the specific capacity of TIL-derived CD8 T-cells to suppress tumor recurrence in resectable HBV-HCC patients"

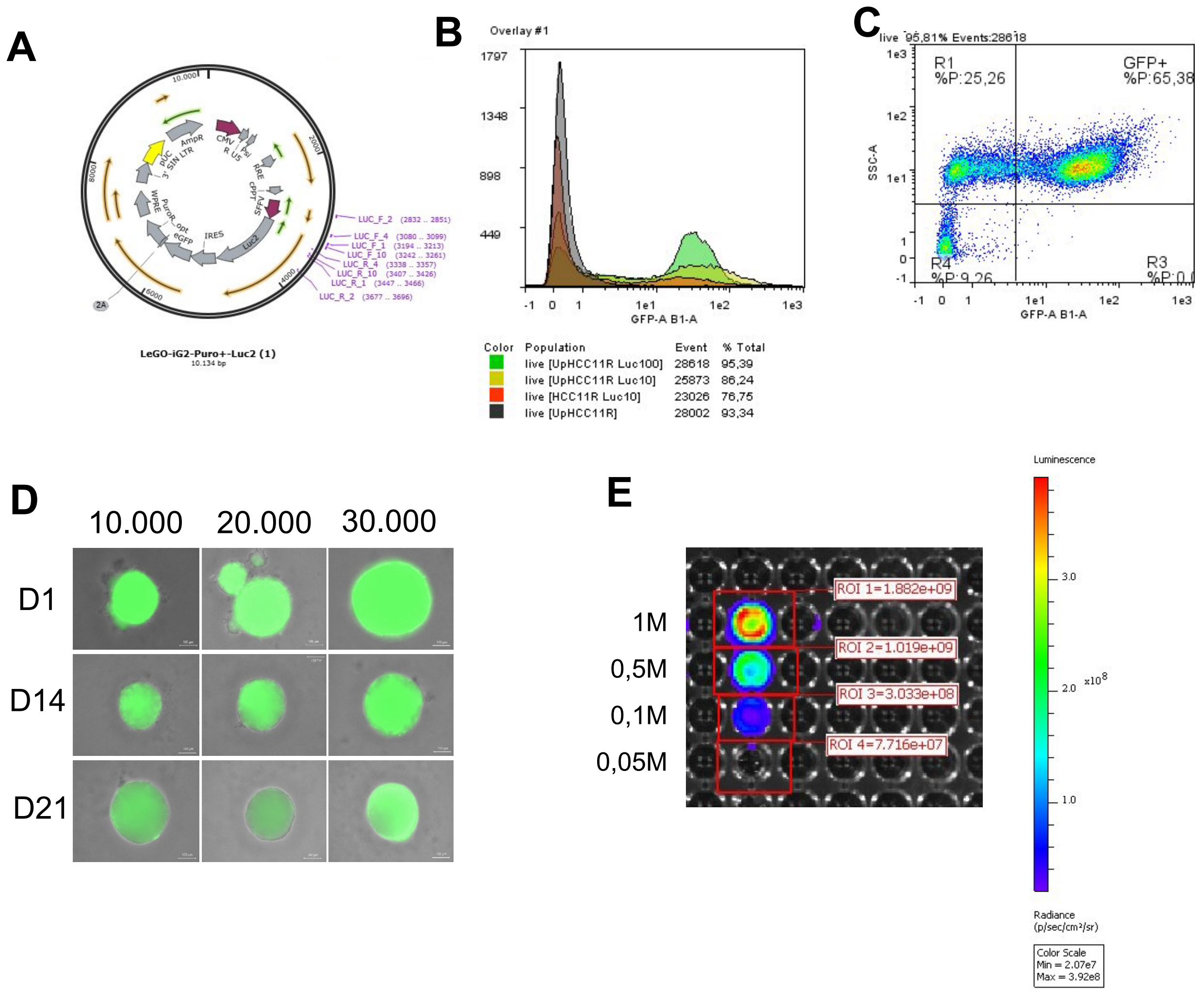

***Supplementary Figure 2:* Characterization of stably transduced generated LC11R_LUC cells, using LeGo-iG2-Puro+-Luc2.**

**Panel A** represents the gene card of the transduced LeGoVector, including the LUC-specific primer pairs. **Panel B** represents histograms overlaid with corresponding events and percentages of GFP-positive populations marked by indicated colour codes. In **panel C,** a representative SSC-A / GFP gate shows the distribution of the LC11R cells 2 weeks after stable transduction and puromycin selection. In panel D, spheroid formation using indicated cell numbers was conducted for 3 weeks and merged captures using bright field, and 488 channels were generated using the Keyence microscope BXZ790. Luciferase production for diluted cell numbers was determined by Luciferin application. The signal was measured and quantified using the In vivo Imaging system in combination with the Living Image Software version 4.7.4 from PerkinElmer, as shown in **panel E.**
