## Supplementary Figure 3 for "Accessing the specific capacity of TIL-derived CD8 T-cells to suppress tumor recurrence in resectable HBV-HCC patients"

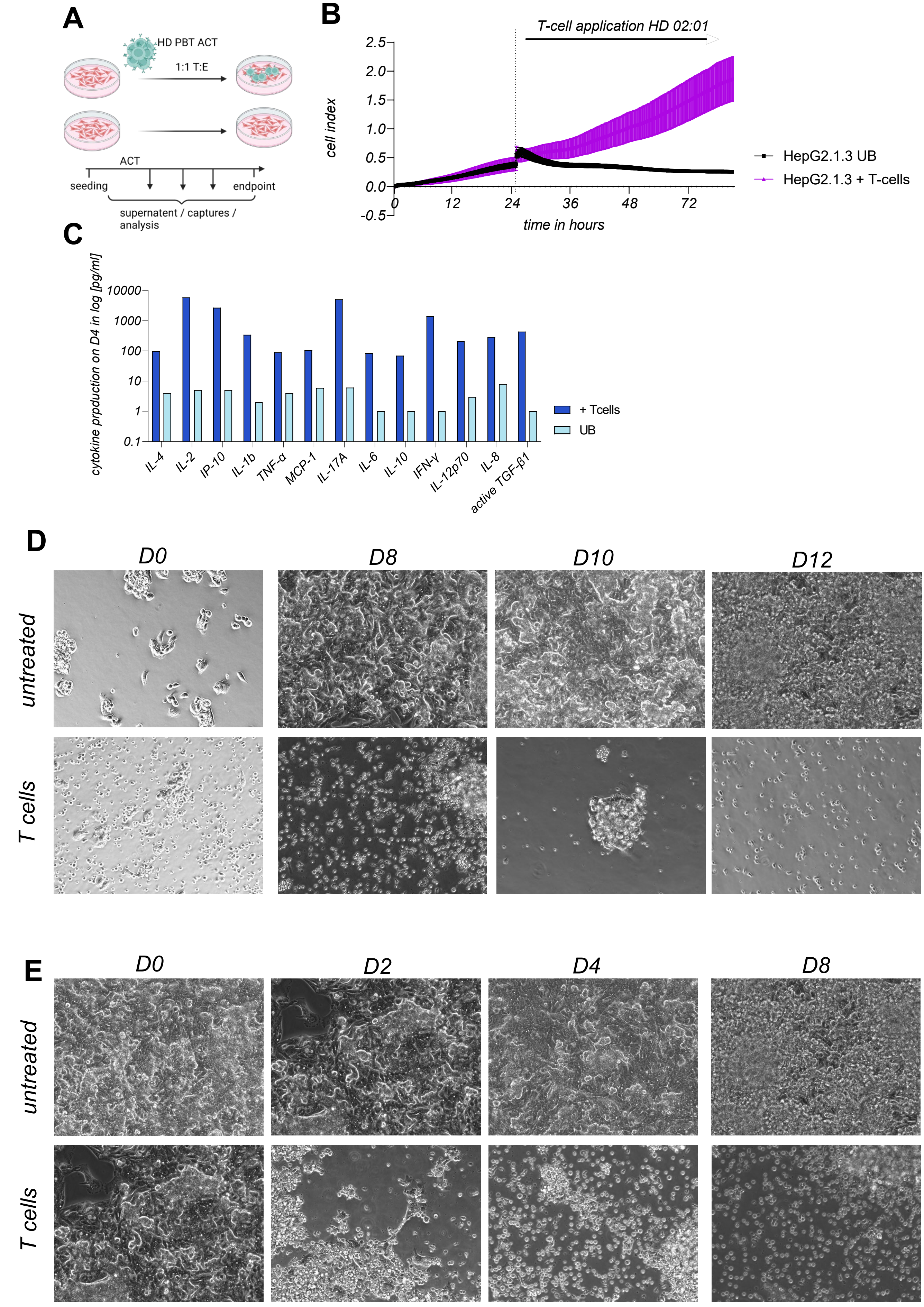

***Supplementary Figure 3:*** **PBT ACT POC for HD02:01 in a matched environment**

**Panel A** represents the experimental setting of the ACT shown in Supplementary Figure 3. In panel **B**, cell index measurement presents the untreated and the treated samples (n = 5) for the indicated time. Panel **C** represents the cytokine measurement after 4 days of PBT ACT in a 1:1 E: T ratio, acquired by multiplex analysis for the treated and the untreated samples (2 replicates from the pooled supernatant). Panel **D** shows representative microcopy captures of the same zone of interest for the indicated time points after ACT, with low cell seeding numbers of LC11R cells and high seeding numbers present in panel **E**, for the indicated time points.
