## Supplementary Figure 5 for "Accessing the specific capacity of TIL-derived CD8 T-cells to suppress tumor recurrence in resectable HBV-HCC patients"

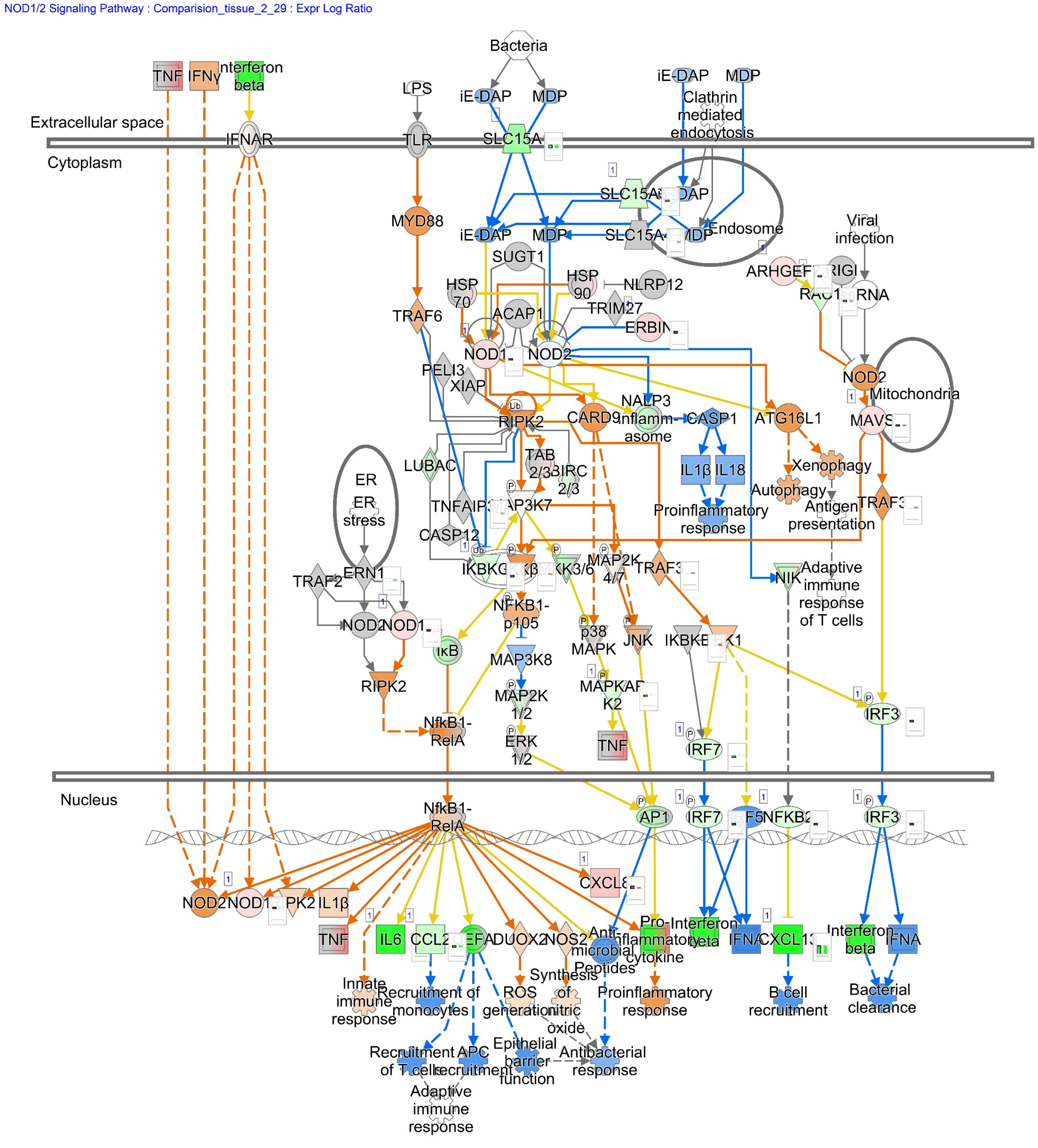


© 2000-2024 QIAGEN. All rights reserved.

***Supplementary Figure 5:*** **NOD1/2 signaling pathway with predicted activation and inactivation for the cancer margin RNAseq analysis.**

DEG from the cancers margin RNA compared to RNA from healthy PHH was used for pathway-related molecule activity and inactivity prediction. Blau indicates the prediction of inactivation, and orange indicates the activation prediction. Green represents downregulation, red represents upregulation, and the shades represent the level differences in the expression changes. Grey arrows and colour represent no differences in the detected DEGs. White tinted parts represent no detection, and yellow arrows indicate a controversial difference between measurement and prediction. Interrupted arrows describe an indirect relation, while solid arrows describe a direct relation. Pathway analysis was performed using IPA software as described in the methods.
