## Supplementary Figure 6 for "Accessing the specific capacity of TIL-derived CD8 T-cells to suppress tumor recurrence in resectable HBV-HCC patients"

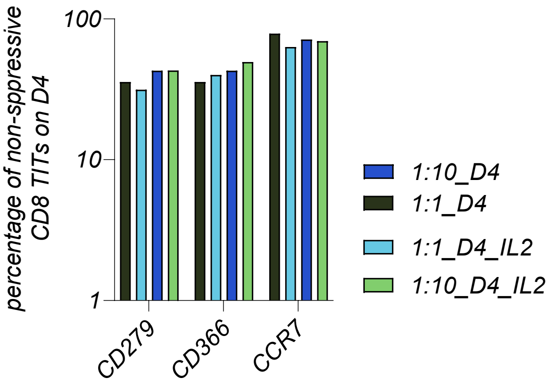

***Supplementary Figure 6:*** **Percentage of non-suppressive CD8 TITs during ACT**

The Figure represents the non-suppressive subpopulation of CD8 T cells derived from patients' tumor-infiltrating lymphocytes, after expansion, stimulation and ACT to the center cell line LC11Z. The data for the indicated groups was determined on day 4 after ACT.
