## Supplementary Figure 7 for "Accessing the specific capacity of TIL-derived CD8 T-cells to suppress tumor recurrence in resectable HBV-HCC patients"

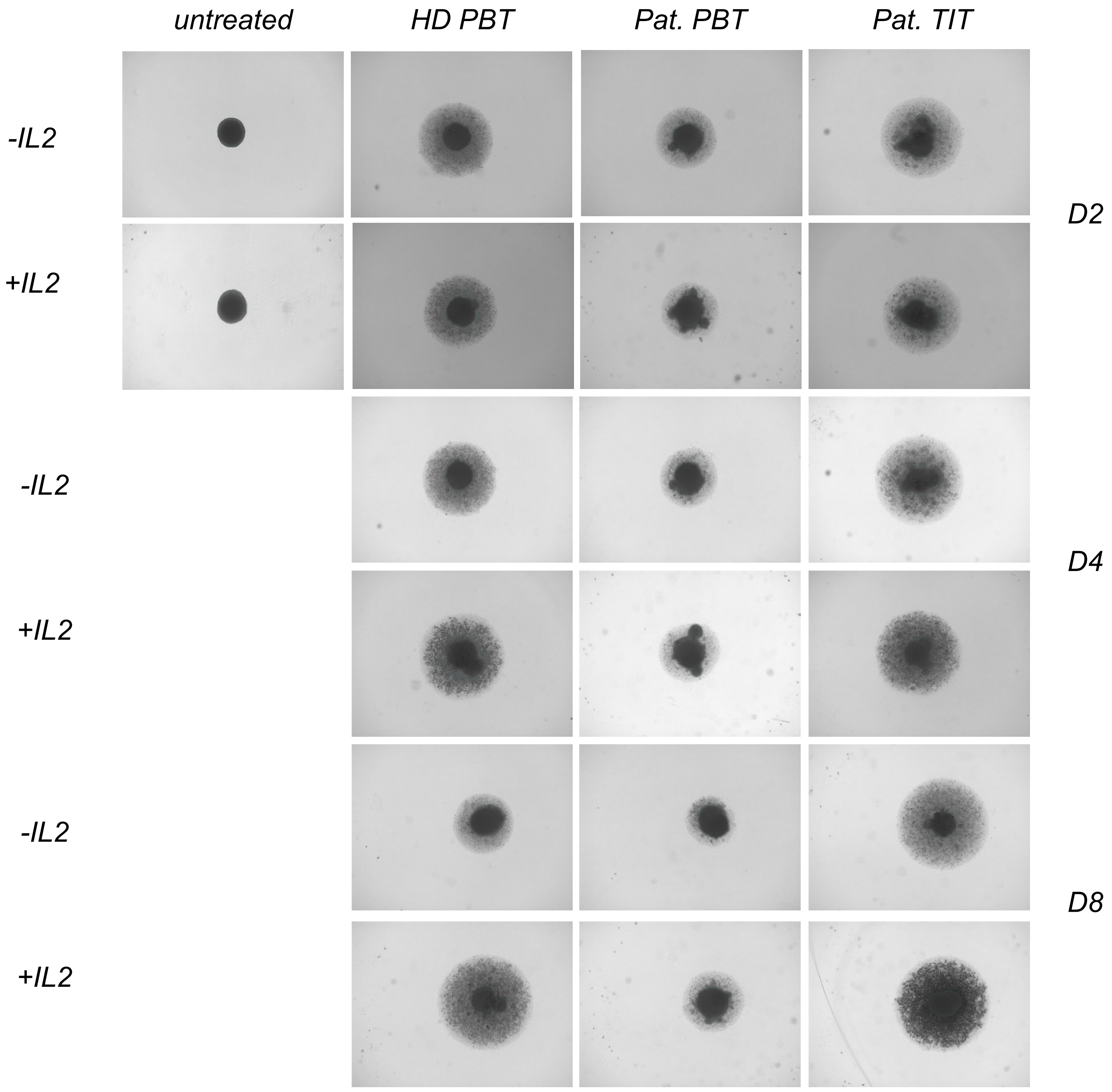

**Supplementary Figure 7: Representative captures following ACT of indicated immune cell types on LC11Z spheroids.**

30.000 cancer center cells (LC11Z) were seeded per U-well and allowed to form spheroids for 3 weeks. After spheroid formation, as illustrated in Figure 4D, ACT of different expanded and stimulated immune cell sources was performed. Spheroids were followed over time, conducting automatic captures using the brightfield channel of the BX-780 Microscope (Keyence, Osaka, Japan). Identical spheroids were used for longitudinal visualization of the treatment, shown in the layout. For each treatment group, n = 3 spheroids were captured and subsequently used for the macro-based automatic area calculation. Areas used for the spheroid size calculation are marked with light green transparent forms, shown in the layout.
