## Supplementary table 1 for "Accessing the specific capacity of TIL-derived CD8 T-cells to suppress tumor recurrence in resectable HBV-HCC patients"

**Supplementary Table 1:** Staining panel for T cell differentiation

| Panel | Marker | Flourochrome |
|---|---|---|
| S10 | CCR7 | BB515 |
| | CD152 | PE |
| | CD272 | BV711 |
| | CD279 | PE-Cy7 |
| | CD3 | PerCP-Cy5.5 |
| | CD4 | APC-H7 |
| | CD45 | BUV805 |
| | CD45RA | BV605 |
| | CD8a | AF700 |
| | TIGIT | V450 |
| S7 | CD103 | BV605 |
| | CD16 | BV421 |
| | CD197 | BB515 |
| | CD20 | PE-CF594 |
| | CD279 | BB790 |
| | CD3 | BB700 |
| | CD366 | BV786 |
| | CD39 | BV650 |
| | CD4 | APC-H7 |
| | CD45 | BUV805 |
| | CD69 | BV510 |
| | CD8a | APC-700 |
