## Supplementary table 2 for "Accessing the specific capacity of TIL-derived CD8 T-cells to suppress tumor recurrence in resectable HBV-HCC patients"

**Supplementary Table 2:** TaqMan Assays used for cDNA amplification

| gene name | Assay-ID |
|---|---|
| HBV S | Pa03453405_s1 |
| Caspase 3 | Hs00234387_m1 |
| IFNgamma | Hs00989291_m1 |
| RPL0 | Hs00420895_gH |
| CCR7 | HS01013469_m1 |
| HNF4A | Hs00230853_m1 |
| HIF1A | Hs00936375_m1 |
| CD44 | Hs01075864_m1 |
| CD8A | Hs00233520_m1 |
| CD4 | Hs01058407_m1 |
