## Supplementary table 3 for "Accessing the specific capacity of TIL-derived CD8 T-cells to suppress tumor recurrence in resectable HBV-HCC patients"

**Supplementary Table 3:** PCR Primer pairs for cf-DNA measurement

| Primer | | Sequence (5'->3') |
|---|---|---|
| LUC_F_1 | Forward primer | CGATATCCGCCACCATGGAA |
| LUC_R_1 | Reverse primer | GCAAGCTATTCTCGCTGCAC |
| LUC_F_2 | Forward primer | AAGTTCAGATCAAGGGCGGG |
| LUC_R_2 | Reverse primer | GCTTTGGAAGCCCTGGTAGT |
| LUC_F_4 | Forward primer | AATCAGCCTGCTTCTCGCTT |
| LUC_R_4 | Reverse primer | GTCCACCTCGATATGTGCGT |
| LUC_F_10 | Forward primer | CGCCATTCTACCCACTCGAA |
| LUC_R_10 | Reverse primer | ATTCAGCCCATAGCGCTTCA |
| ALU_F_1 | Forward primer | CCCGAGTAGCTGGGATTACA |
| ALU_R_1 | Reverse primer | CCCGAGTAGCTGGGATTACA |
| MTCO_F_1 | Forward primer | TAAACTTCAACCAACACCGT |
| MTCO_R_1 | Reverse primer | TAGACTTCTGGGTGGCCAAAGA |
| MTCO_F_2 | Forward primer | GACCTGATGCACTGAGGTTT |
| MTCO_R_2 | Reverse primer | GTTTACGAGGCTTCTTCTG |
